# T cell ectosomes promote antibody responses through cognate TCR-pMHC interactions

**DOI:** 10.1101/2025.07.30.667776

**Authors:** Fenglei Li, Benjamin Schmitz, Henriette A. Remer, Julia Brasch, Wesley I. Sundquist, Kaushik Choudhuri

## Abstract

Protective antibody-mediated immunity requires effective T cell-mediated help. Recognition of peptide antigens presented by major histocompatibility complex class II molecules (pMHCII), *via* cognate T cell antigen receptors (TCR), activates CD4^+^ T helper cells to upregulate expression of CD40L and helper cytokines. CD40L-CD40 interactions at T-B cell synapses and secreted cytokines are well established mediators of T cell help. Whether engaged pMHCII also transmits signals that help B cells remains unresolved. Here, we show that TCR-enriched nanoscale vesicles shed by activated T cells (ectosomes) are transferred to antigen-primed B cells, where they engage and cluster cognate pMHCII, triggering signaling and specific IgG antibody production. Disruption of ectosome release attenuates B cell antibody production, while native and synthetic ectosomes boost antibody responses. We conclude that T cell ectosomes constitute a new modality of help for B cells, delivered through engagement of pMHCII by cognate ectosomal TCR.

---

Collaboration between T cells and B cells is fundamental to adaptive immunity and the production of protective antibodies against infectious pathogens or following vaccination (*1*). Binding and internalization of cognate antigens by surface-expressed B cell receptors (BCR) activates B cells to upregulate expression of CD69 (*2*), major histocompatibility class II molecules (MHCII) (*3*), and costimulatory receptors such as CD80/86 (*4*). BCR-bound protein antigens are rapidly internalized, processed into peptide fragments, and displayed on the B cell surface in complex with MHC class II molecules (pMHCII) (*3*). Recognition of pMHCII, *via* surface-expressed cognate T cell antigen receptors (TCR), activates T helper cells to upregulate expression of CD40 ligand (CD40L) and helper cytokines including IL-4 and IL-21(*5*). Activated CD4^+^ T helper cells provide contact-dependent ‘help’ to B cells through CD40L-CD40 interactions at T-B cell synapses (*6*), and by secretion of soluble helper cytokines (*5, 7*), that together promote B cell survival, proliferation, differentiation and isotype class-switched antibody production.

While recognition of pMHCII displayed on B cells activates cognate T helper cells during T-B cell collaboration (*6, 8*), the role of pMHCII engagement in B cell responses remains unclear. Artificial cross-linking of MHCII by antibodies triggers intracellular signaling (*9–11*) in IL-4- primed B cells, while engagement of pMHCII with bead-immobilized cognate TCRs induces phosphorylation of BCR-associated signaling subunits CD79A/B and activation of the B cell tyrosine kinase Syk (*12*). This results in intracellular Ca^2+^ influx and non-specific antibody production *in vitro* (*12–14*). Guy *et al* (*15, 16*) observed that antigen-activated T helper 2 (Th2)- polarized T cell clones release large aggregates of ‘cell-free’ TCRs into culture supernatants, which stimulated antibody production in cocultures with syngeneic memory-enriched splenic B cells (MEBC), isolated from mice immunized with haptenated protein antigens recognized by the Th2 cell clones. Notably, antibody production was not induced MHC-mismatched T cell clones, demonstrating that TCR-induced antibody production was MHC-restricted. However, no physiological mechanism is known for shedding of membrane-tethered TCRs, or for their multimerization in solution, calling into question their relevance to physiological B cell responses. We have shown that activated CD4^+^ effector T cells release TCR-enriched nanoscale vesicles (ectosomes) from their surface in an ESCRT (<u>e</u>ndosomal <u>s</u>orting <u>c</u>omplexes required for transport)-dependent manner (*17*), although their precise functions remain unclear. CD4^+^ T effector cell-derived ectosomes have been shown to activate dendritic cells *in vitro* through non-antigen-specific CD40L-mediated engagement (*18*), but TCR-mediated functions remain unexplored. Since T cell ectosomes typically display hundreds of binding-competent TCRs on their surface (*17*), we investigated whether ectosomes produced by differentiated helper T cells can function as a third modality of ‘help’ for B cell antibody production, distinct from direct contact-dependent T-B cell interactions and soluble signals.

## Extracellular TCR is packaged in T cell ectosomes

We first established a scalable protocol for the bulk production and isolation of T cell ectosomes with a uniform TCR clonotype. For this, we chose *in vitro*-Th2 polarized CD4^+^ OTII TCR transgenic T cells (*19*), which recognize ovalbumin peptide OVA^323-339^ restricted by murine MHC class II molecule I-A^b^ (pOVA/I-A^b^) (fig. S1A). Notably, *in vitro*-differentiated OTII Th2 cells, which upregulated GATA-3 (fig. S1B) and secreted IL-4 when activated by antigen-presenting B cells (*5*) (fig. S1C), produced two-fold more ectosomes than undifferentiated Th0 effector cells, as measured by nanoparticle tracking analysis (NTA) (fig. S1D). OTII Th2 cells were also superior in providing help for isotype class-switched NP-specific IgG antibody production in coculture with nitrophenol (NP)-specific transgenic QM B cells (*20*) primed with NP-haptenated ovalbumin (NP-OVA) (fig. S1E).

Expanded quiescent OT-II Th2 cells were purified and re-stimulated with plate-adsorbed anti-CD3/CD28 antibodies to elicit ectosome release into exosome-depleted culture supernatants (Fig. 1A and fig. S1A), and isolated using standard ultra/centrifugation steps (fig. S1A) (*21*). Resting T cells released few ectosomes, but polarized stimulation with solid phase anti-CD3/CD28 antibodies increased production by >10-fold after 48 hours of stimulation, yielding ∼10^9^ ectosomes/million T cells (fig. S2A) with a mean diameter of ∼140nm (Fig 1,B and C and fig. S2B). These were comparable in number and size to ectosomes elicited using *bona fide* pOVA/I- A^b^ antigen complexes and CD80 (fig. S2, C to E), and contained similar levels of TCR-ζ, CD3ε and TSG101 (fig. S2F).

**Fig. 1.**
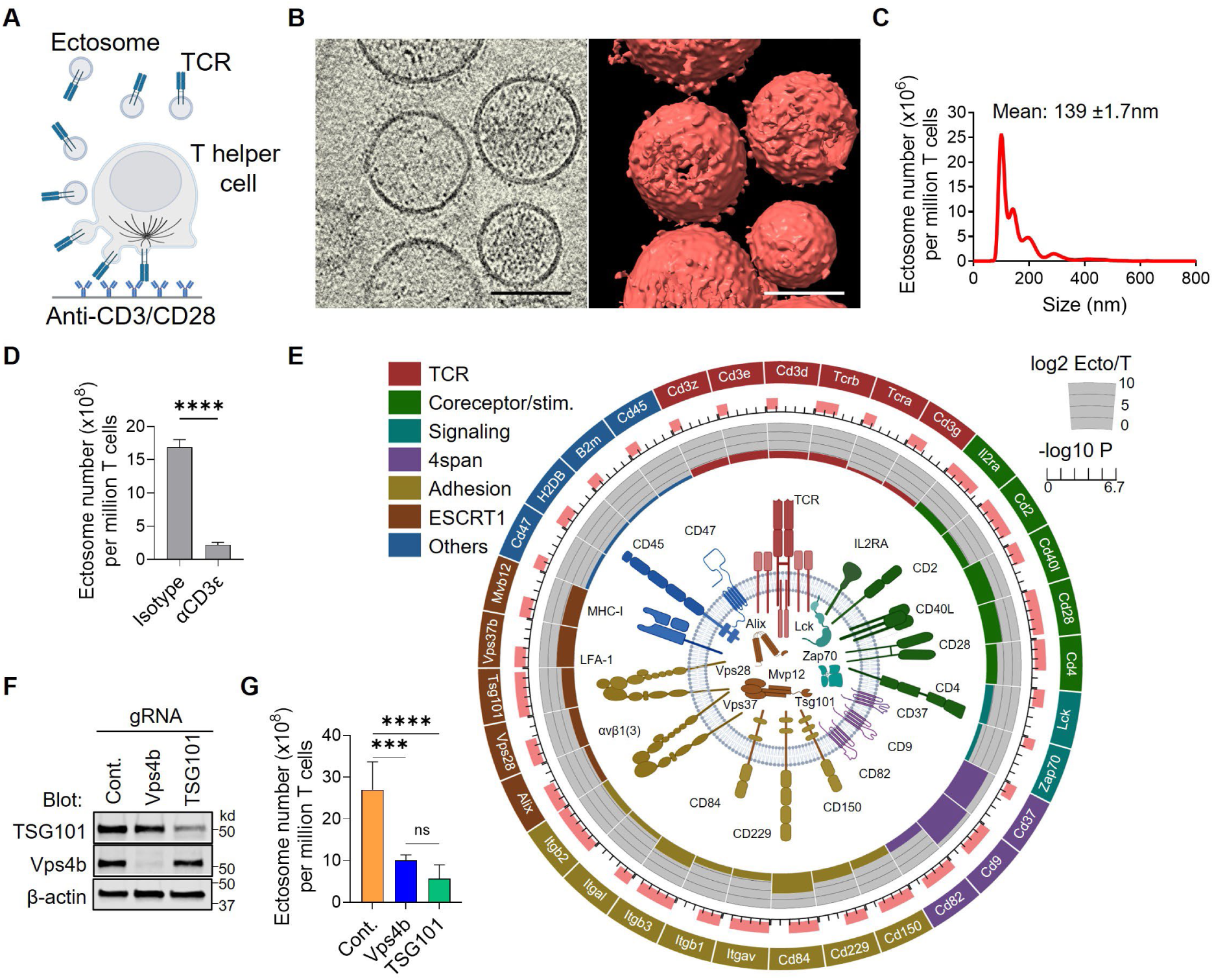
Ultrastructure, proteomics and biogenesis of OTII T helper cell-derived ectosomes. (**A**) Schematic of TCR-enriched ectosomes released from *in vitro*-differentiated OTII T helper 2 (Th2) cells upon activation with solid phase immobilized anti-CD3/CD28 antibodies. Ectosomes were isolated by ultracentrifugation. (**B**) (Left panel) Middle slice of segmented tomogram of T cell ectosomes in vitrified ice, acquired by electron cryotomography (ECT). (Right panel) Isosurface rendering of ECT reconstructions of T cell ectosomes. Scale bar, 100nm. (**C**) NTA quantitation of T cell ectosome size. Mean ± s.e.m. is indicated; *n* = 20,000 trackable particles. (**D**) Depletion of isolated ectosomes in suspension using bead-immobilized anti-CD3ε antibody or a bead-immobilized non-specific isotype control antibody, measured by NTA. Data are mean ± s.e.m. compared using two-tailed *t*-test. Results are representative of 4 independent experiments. (**E**) Circular graph showing selected proteins enriched in ectosomes grouped and color-coded according to function. Color-coded bars in grey calibrated plot depict log2 transform of enrichment ratio relative to parental Th2 cells, pink bars depict -log10 transformed P values. (Central inset) Cartoon of an OTII Th2 cell-derived ectosome based on the corresponding proteomic analysis, depicting membrane-tethered surface receptors and luminal ESCRTs and kinases, color coded to correspond to groupings in proteomic analysis. (**F**) Immunoblot of CRISPR-mediated gene knockout (KO) of ESCRT members TSG101 and VPS4b in OT-II^CAS9^ T cells. Cont., non-targeting gRNA transduced control. (**G**) NTA quantitation of ectosome release from TSG101 and VPS4 KO OT-II^CAS9^ T cells following activation with plate-bound anti-CD3 antibody. Data are representative of 4 independent experiments. Cont., non-targeting gRNA transduced control. Means were compared by one-way ANOVA corrected for multiple comparisons. ****, P < 0.0001; ***, P < 0.001; ns, not significant (P > 0.05).

Cryoelectron microscopy and electron cryotomography of isolated ectosomes in vitrified ice established that T cell ectosomes exhibited typical vesicular morphology, and confirmed their increased production following T cell activation (Fig. 1B and fig. S2, G to J). Segmented tomograms and rendered three-dimensional models (Fig. 1B) revealed discrete spherical vesicles of ∼100-150 nm in diameter (fig. S2J and Movie S1), which were bounded by well-defined lipid bilayers (Fig. 1B). Small particle flow-cytometry of ectosomes, labelled with green fluorescent lipophilic dye PKH67 for gating, and anti-TCRβ and -CD40L antibodies, revealed a single relatively homogeneous TCRβ^+^CD40L^+^ population (fig. S2, K and L). In agreement with this, >85% of ectosomes contained TCR on their surface, as determined by immunodepletion with anti- CD3-conjugated ferromagnetic beads (Fig. 1D), demonstrating that TCR is shed by activated OTII T cells as TCR^+^ ectosomes.

## Ectosomes concentrate effectors of help for B cells

Polarized stimulation of T cells with solid phase anti-CD3ε antibodies enhanced TCR incorporation into ectosomes relative to stimulation with saturating concentrations of soluble antibodies, or with soluble mitogenic PMA/ionomycin stimuli (fig. S2M), suggesting that polarized TCR engagement promotes TCR sorting into ectosomes. In contrast, incorporation of cytoplasmic TSG101 was not sensitive to TCR-engagement, while β-actin was uniformly excluded from ectosomes, pointing to distinct sorting mechanisms that control ectosome surface and cargo composition (fig. S2M).

To characterize the proteome of OTII T cell ectosomes more comprehensively, we performed LC-MS mass spectrometry (*22*) (Fig. 1E and fig. S3). This confirmed enrichment of the multisubunit TCR, along with its proximal signaling kinases Lck and ZAP-70, consistent with their incorporation into ectosomes as a multimolecular post-activation complex (*17, 23, 24*). Importantly, CD40L was enriched by ∼40-fold in ectosomes relative to parental T cells, in keeping with a high capacity for providing help to B cells. Several ESCRT complex proteins, including the key proximal mediators of vesicle budding, ALIX (*25*) and the heterotetrameric ESCRT-I complex (TSG101, VPS28, Vps37b, and Mvb12a) (*26*) (Fig. 1E and fig.S3A), were also enriched in ectosomes, as were cell surface-expressed nutrient transporters and ion transport channels (fig. S3, B and C), reflecting the plasma membrane origin of T cell ectosomes. Consistent with this, proteins associated with endomembranes of the Golgi apparatus, endoplasmic reticulum, nucleus and mitochondria were under-represented in ectosomes (fig. S3, B and D). Key adhesion and costimulatory receptors that promote binding to the B cell surface were also concentrated in ectosomes (Fig. 1E). This included enrichment of the T cell coreceptor CD4 (*27*), which binds to MHCII in *trans*, along with SLAM family adhesion receptors and integrins known to be important for adhesion to B cells and humoral responses (*28–31*).

## ESCRT-dependence of T cell ectosome biogenesis

We have shown that transfer of TCR to B cells can be disrupted by suppressing expression of proteins in the ESCRT pathway that are involved in budding and release of vesicles from the plasma membrane (*17*), although the contribution of ESCRT proteins to ectosome production and function in differentiated T helper cells has not been examined. We therefore employed CRISPR editing to knockout TSG101, an ESCRT I protein that initiates HIV/ectosome budding from the plasma membrane (*32*), and VPS4B, a subunit of the VPS4 AAA+-ATPase which catalyzes the final step of membrane scission that results in enveloped virus/ectosome release (*32*). Lentiviral transduction of Th2-polarized cells, harvested from F1 progeny of crossed OTII and Cas9 transgenic mice (OTII^CAS9^) (*33*), with expression vectors for gRNAs targeting TSG101 and VPS4B (Table S2), resulted in ∼90-95% reduction in their respective protein expression levels (OTII^CAS9^ KO T cells) (Fig. 1F). Deletion of ESCRTs did not affect differentiation into Th2 cells as indicated by effective GATA3 upregulation (fig. S4A), or secretion of IL-4 (fig. S4B). Surface TCR levels (fig. S4C) and TCR-mediated activation, as measured by CD69 upregulation (fig. S4, D and E) and secretion of IL-2 (fig. S4F), were also unaffected in OTII^CAS9^ KO T cells. In contrast deletion of TSG101 or VPS4B significantly reduced vesicle release by activated OTII^CAS9^ T cells by ∼3-5 fold compared to T cells transduced with a non-targeting gRNA expression vector (Fig. 1G), highlighting their central role in the production and release of ectosome from T cells.

## T cell ectosomes induce antigen-specific signaling and antibody production in B cells

We reasoned that binding of multimeric ectosome-associated TCRs to cognate pMHCII complexes could potentially deliver high avidity antigen-specific signals to antigen-primed B cells. To test this, we first compared inducible protein tyrosine phosphorylation in NP-OVA-activated and IL-4-primed B cells following incubation with OTII Th2 cell-derived ectosomes (Fig. 2, A and B). Incubation of primed NP-specific BCR-transgenic QM B cells (fig. S6A) with ectosomes for 2 minutes induced several phosphorylated protein bands (Fig. 2B), one of which was the protein tyrosine kinase Syk (Fig. 2B), a critical proximal effector of B cell signaling downstream of the BCR known to associate with MHCII in B cells (*12*), including activation of PLCγ (*34*), which drives intracellular Ca^2+^-signaling. Antibody-mediated MHCII-crosslinking resulted in a similar pattern of protein tyrosine phosphorylation in QM B cells (Fig. 2B) suggesting that MHCII clustering is a necessary step for signal transduction. To establish whether T cell ectosomes induced downstream signaling in B cells, we used flow-cytometry to monitor Ca^2+^ signaling in QM B cells in response to T cell ectosomes. B cells were primed with IL-4 and NP-OVA or noncognate NP-HEL, and loaded with the ratiometric fluorescent Ca^2+^ indicator Indo-1(*35*). Ectosomes-induced B cell signaling was strictly antigen-specific as incubation with OTII T cell-derived ectosomes triggered rapid Ca^2+^-influx in primed B cells loaded with NP-OVA, detected as increased Indo-1 420/475nm fluorescence ratio, while no increase was detected in B cells loaded with the NP-HEL (Fig. 2C). To test ectosome-mediated antibody responses in more physiological polyclonal B cells, we hyperimmunized C57bl/6 mice with multiple rounds of NP-BSA inoculation (Fig. 2D), to generate polyclonal NP-specific primary memory-enriched B cells (MEBC) (*36*) (fig. S5A and B). NP-specific splenic B cells from hyperimmunized mice were enriched in IgD^-^ /IgG^+^ B cells (Fig. 2, E and F), and exhibited a memory phenotype, with upregulated surface expression of CD11a, CD81, CD180, CD274 (PD-L1) and CD205 (fig. S5, A and B) (*36*). Surprisingly, we also observed the appearance of TCR^+^CD40L^+^extracellular vesicles in the plasma fraction of blood collected 1 week after the first vaccination, which increased incrementally upon each subsequent inoculation (fig. S5C), suggesting that T cell activation during immunization results in T cell ectosome release that is detectable in the circulation.

**Fig. 2.**
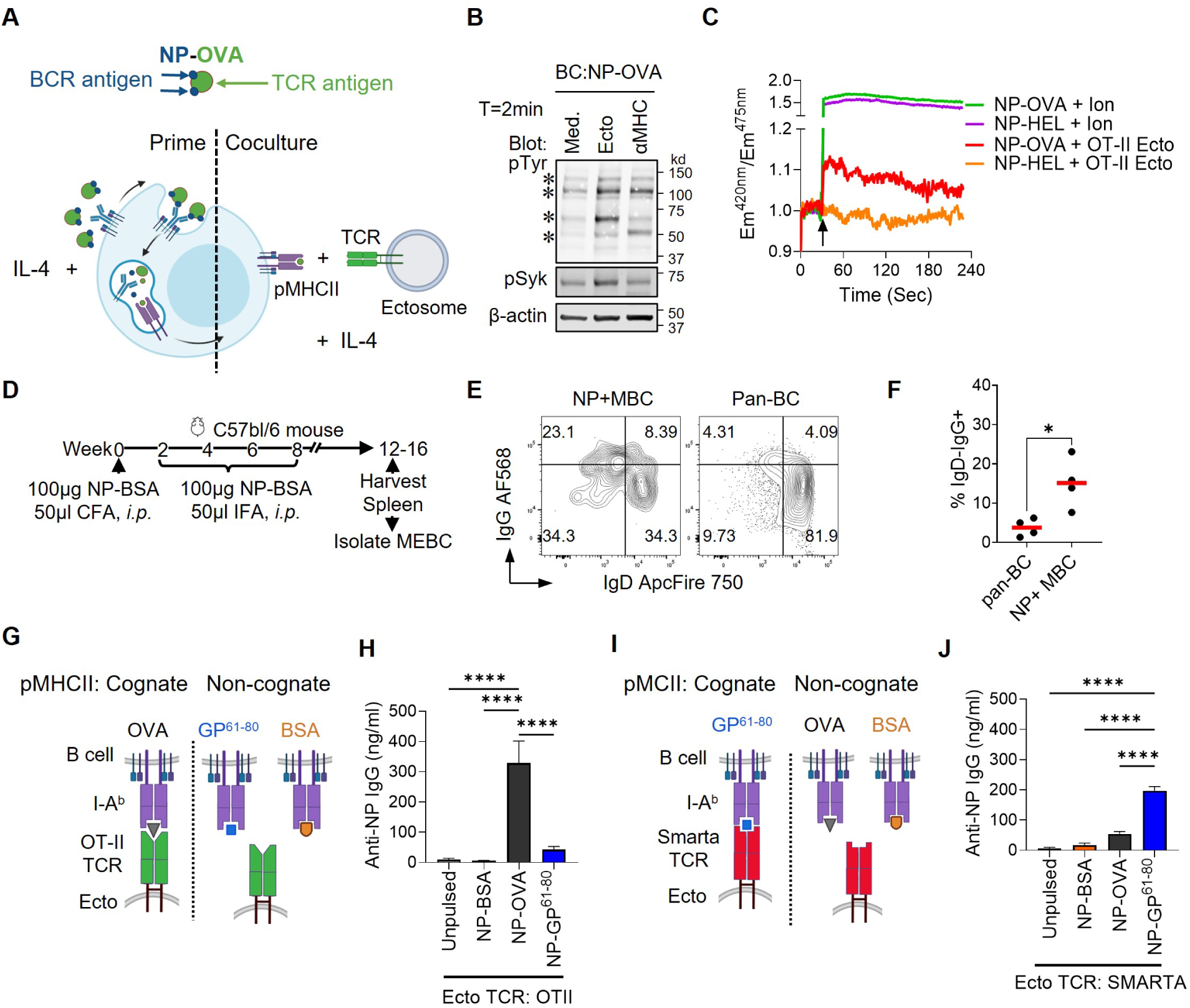
T cell ectosomes induce antigen-specific signaling and antibody production in B cells. (**A**) Schematic of experimental format. (Left side of dashed line) Depicts IL-4 priming of NP-specific QM B cell and activation with NP-conjugated ovalbumin antigen (NP-OVA), (Right side of dashed line) Depicts cognate TCR-bearing ectosome and QM B cell displaying pMHCII cocultured in the presence of IL-4. (**B**) Primed B cell displaying pOVA/I-A^b^ pMHCII complexes cocultured with ectosomes and IL-4. Immunoblot of tyrosine phosphorylated proteins (pTyr) and Syk Tyr352 in NP-OVA-loaded QM B cells, unstimulated (Med.) or treated with OTII-derived ectosomes (Ecto), or anti-MHC II-antibody (αMHC). Results are representative of 4 independent experiments. Asterisks indicate positions of phosphorylated protein bands. QM B cells (BC) were activated with NP-OVA. (**C**) Intracellular Ca^2+^-influx in QM B cells pulsed with NP-OVA or NP- HEL antigens following addition of ectosomes (arrow) from activated OT-II T cells (Ecto). The ratio of Indo-1 fluorescence at 420nm (Ca^2+^-bound) and 475nm (Ca^2+^-unbound) was taken as measure of intracellular Ca^2+^ levels. Ion, cells were treated with 10µM ionomycin as a positive control; results are representative of 3 independent experiments. (**D**) Timeline of protocol for hyperimmunization of C57bl/6 mouse with NP-BSA for isolation of splenic memory-enriched B cells (MEBC). (**E**) Contour plots of surface IgD and IgG levels on on NP^+^MBC (CD19^+^CD38^+^CD81^+^NP^+^ live singlets) and pan-B cells (CD19^+^ live singlets) populations measured by flow cytometry. (**F**) Quantitation of the percentage of class-switched IgD-/IgG+ B cells in the indicated B cell populations. Data points represent individual mice. Horizontal red lines indicate means. Means were compared by two-tailed *t*-test. (**G**) and (**I**) Schematic of TCR and antigen combinations used to measure the antigen-specificity of MEBC antibody production in the assay outlined in *(A)*. (**H**) and (**J**) Quantitation of antibody production by MEBC primed with the indicated NP-haptenated protein antigens and cocultured with ectosomes expressing the indicated TCR. Data are means ± s.d.. Means were compared using 1-way ANOVA corrected for multiple comparisons. . ****, P < 0.0001; P < 0.05.

To determine whether T cell ectosomes alone can induce antibody production in B cells, we activated MEBC with NP-BSA, NP-OVA or a NP-conjugated synthetic antigen comprising a fusion of GFP and the LCMV-derived gp61-80 peptide (NP-GFP.GP^61-80^) (*37*), which is an I-A^b^- restricted epitope recognized by SMARTA-1 TCR-transgenic CD4+ T cells (Fig. 2, A and G and fig. S10A). All three NP-conjugated antigens elicited similar levels of CD69 upregulation when incubated with NP-specific QM B cells, indicating comparable BCR-mediated activation potency (fig. S6, B and C). Ectosomes derived from Th2-polarized OTII T cells induced robust NP-specific class-switched IgG antibody secretion by IL-4-primed and NP-OVA-activated MEBC (Fig. 2H). Antibody production was highly antigen-specific, as MEBC activated with NP-BSA or NP- GFP.GP^61-80^ failed to produce NP-specific antibodies in response to OTII-T cell-derived ectosomes (Fig. 2, G and H). Conversely, ectosomes derived from Th2-polarized SMARTA T cells induced robust NP-specific antibody production in IL-4-primed and NP-GFP.GP^61-80^- activated MEBC, while antibody production was substantially attenuated in B cells activated with non-specific antigens NP-BSA or NP-OVA (Fig. 2, I and J). Together these results demonstrate that TCR engagement of cognate pMHCII on B cells is necessary for T cell ectosome-mediated signaling and class-switched antibody production.

## TCR transfer to B cells clusters pMHCII and boosts antibody production

To investigate molecular trafficking of TCR and MHCII during T-B cell interactions *in vitro*, we used flow-cytometry to monitor TCR and I-A^b^ transfer in cocultures of NP-OVA-activated QM B cells and OTII Th2 cells (Fig. 3A and fig. S6A). QM B cells activated with NP- conjugated hen egg lysozyme (NP-HEL) were used as a specificity control. NP-OVA and NP- HEL elicited comparable B cell activation, as measured by CD69 upregulation (fig. S6, B and C).

**Fig. 3.**
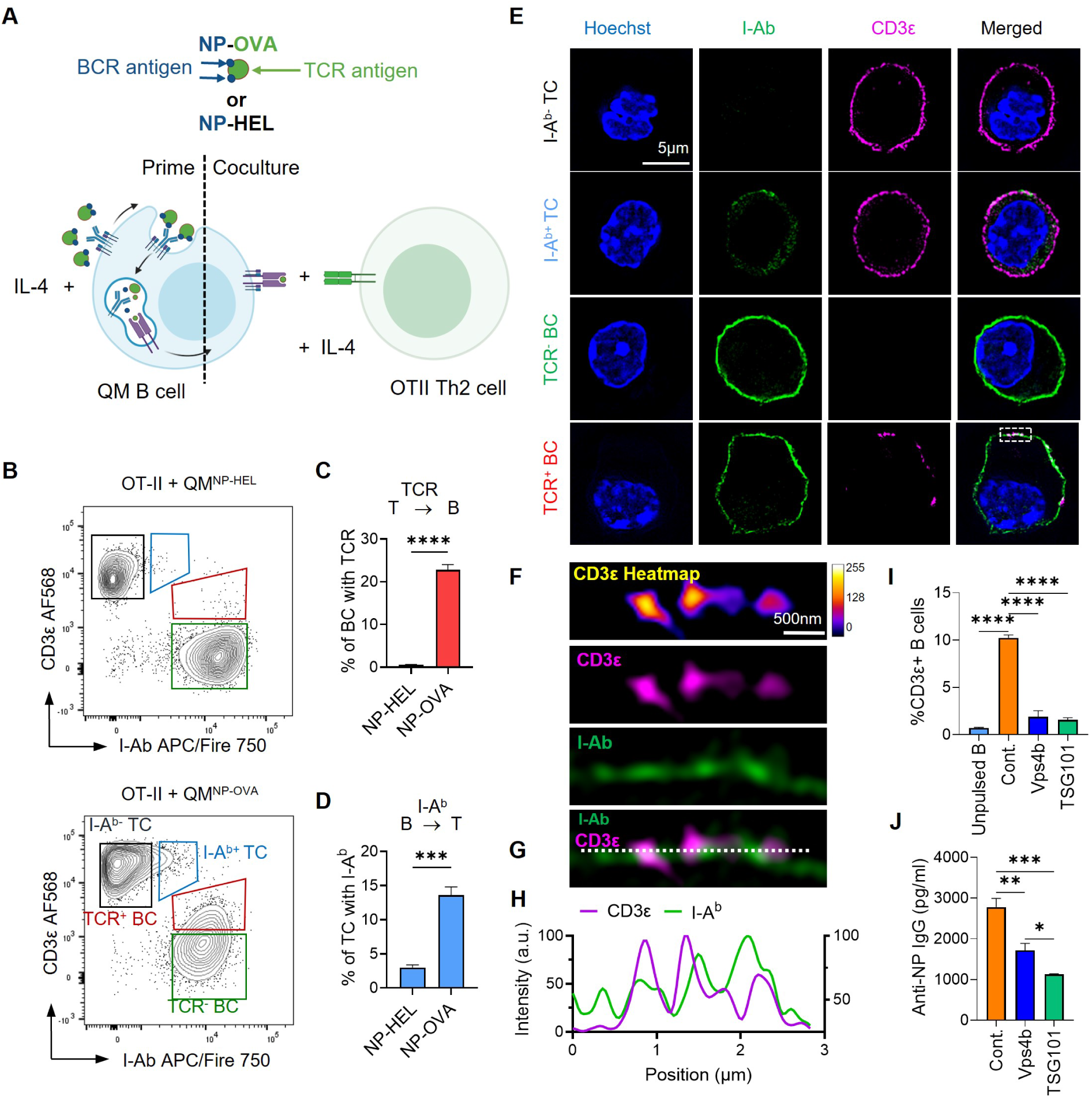
ESCRT-dependent TCR transfer delivers help for B cell antibody production. (**A**) Schematic of experimental format of T-B coculture. (**B**) Flow-cytometry analysis of QM B cells pulsed overnight with 200μg/ml NP-OVA or NP-HEL antigens and co-cultured with Th2 OT-II T cells for 24 hrs. Cells were then PFA-fixed and stained with fluorescently labeled anti-I-A^b^ antibody (upregulated in activated B cells) and anti-CD3ε antibody (TCR and T cell marker). Regions: black square: T cells (I-A^b-^ TC), green square: B cells (TCR^-^ BC), red polygon: TCR transfer to B cells (TCR^+^ BC), cyan polygon: I-A^b^ transfer to T cells (I-A^b+^ TC). Quantitation of percentage of (**C**) B cells with transferred TCR and (**D**) T cells with transferred I-A^b^, in T and B cell cocultures. Mean ± s.d. is shown, results are representative of 4 independent experiments. Means were compared using two tailed *t*-test. (**E**) Representative structured illumination microscopy (SIM) images of T and B cells sorted following coculture using the indicated gates. PFA-fixed cells were permeabilized and labeled with the indicated antibodies conjugated to photostable fluorescent dyes and post-fixed with glutaraldehyde prior to sorting and imaging. (**F)** Magnified region of sorted B cell corresponding to white dashed and boxed region in (E). (**G**) Overlay of TCR and I-A^b^ fluorescence shown in (F). (**H**) Fluorescence intensity of TCR (CD3ε) and I-A^b^ on the B cell surface, measured along the white dotted line in (G). a.u., arbitrary units. (**I**) Quantitation of TCR transfer to NP-OVA-primed QM B cells in cocultures with the indicated ESCRT KO OTII^CAS9^ T cells. Results are representative of 4 independent experiments. Mean ± s.d. is shown. Means were compared by 1-way ANOVA corrected for multiple comparisons. (**J**) Quantitation of IgG antibody production by NP-OVA primed by QM B cells, as measured by ELISA of culture supernatants after 7 days coculture with the indicated ESCRT KO OTII^CAS9^ T cells. Results are representative of 4 independent experiments. Mean ± s.d. is shown. Means were compared by 1-way ANOVA corrected for multiple comparisons. ****, P < 0.0001; ***, P < 0.001; **, P < 0.01; *, P < 0.05.

Transfer of TCR to B cells peaked at 24hrs coculture (fig. S6, D and E), and was strictly antigen-dependent, with ∼20% of NP-OVA activated B cells containing transferred TCR, while TCR transfer was negligible in cocultures with B cells activated with NP-HEL (Fig. 3, B and C). Acquisition of B cell-expressed I-A^b^ by T cells was also apparent, although somewhat less antigen-specific than TCR transfer to B cells (Fig. 3, B and D). To determine the subcellular location and distribution of transferred TCR and I-A^b^, antibody-labelled T and B cells were isolated by cell-sorting of T-B cell cocultures and labelled with antibodies against TCR and I-A^b^, conjugated with photostable fluorescent dyes and imaged by super-resolved structured illumination microscopy (SIM). Four cell populations were sorted and imaged by SIM to determine the distribution of bidirectional molecular transfer: TCR^+^ or TCR^-^ B cells and I-Ab^+^ or I-A^b-^ T cells (Fig. 3,B and E). Although a few puncta of B cell-derived I-A^b^ acquired by T cells (I-Ab^+^ TC) were present on the cell surface (murine T cells do not express MHCII), it was predominantly located in diffuse cytoplasmic compartments (Fig. 3E). In contrast, transferred TCR (TCR^+^ BC) was only detected as bright puncta at the B cell surface (Fig. 3E) colocalized with accumulated of I-A^b^ (Fig. 3, G and H), consistent with multivalent binding and clustering of pMHCII by ectosome-tethered OTII TCRs.

To determine whether the TCR-containing ectosomes that are transferred to B cells provided help for antibody production, we next cocultured NP-OVA-loaded QM B cells with OTII^CAS9^ T KO cells. Deletion of TSG101 or VPS4B resulted in marked reduction of TCR transfer to co-cultured antigen-activated QM B cells (Fig. 3I), consistent with abrogation of intercellular transfer of TCR-enriched ectosomes. Coculture of TSG101- or VPS4B-deleted OTII^CAS9^ T cells resulted in a ∼50% reduction in NP-specific antibody production by NP-OVA activated QM B cells in T-B cell (Fig. 3J), suggesting that in *in vitro* cocultures, ectosome-mediated help was approximately equal in magnitude to the help provided during direct T-B cell interactions for B cell antibody production.

## *In vivo* TCR transfer to B cells and antibody responses

We next investigated whether the transfer of TCR-enriched ectosomes that we observed *in vitro* could also be detected in secondary lymphoid organs during an antigen-induced polyclonal immune response *in vivo*. For this, we seeded C57bl/6 mice intravenously with 10^6^ *in vitro*- expanded OTII Th2 cells to increase the precursor frequency of I-A^b^/OVA-specific T helper cells, and inoculated mice intraperitoneally with NP-OVA in complete Freund’s adjuvant one day after seeding (Fig. 4A). To allow sufficient time for germinal center reactions to develop, we harvested spleens 7 days post-immunization (*38*) for immunophenotypic analysis by flow-cytometry (Fig. 4A). Within the live T cell compartment (fig. S7A), we detected robust expansion of I-A^b^/OVA- specific CD4^+^ T cells using a fluorescently-tagged I-A^b^/OVA tetramer (Fig. 4B and fig. S7, B and C), of which ∼30% were CXCR5^+^PD-1^+^Bcl6^+^ICOS^hi^CD40L^hi^ T follicular helper cells (*39*) (Fig. 4C and fig. S7, D to F), which specialize in providing help to germinal center B cells (*40*). Transfer of TCR to splenic B cells was readily detectable (Fig. 4 D and E), and was substantial skewed toward GL7^hi^/Fas^+^ B cells (*41, 42*), suggesting that TCR-mediated ectosomal help is preferentially associated with responding, and presumably antigen-specific, germinal center B cells (Fig. 3G). In support of this, B cells with transferred TCR also accumulated markedly more mutations in their BCR VH-domains, especially within complementarity determining regions (CDR) (Fig. 3J), in keeping with the notion that transferred ectosomal TCR engages cognate pMHCII on B cells, triggering signals that contribute to central downstream processes, including the initiation of somatic hypermutation (*43*).

**Fig. 4.**
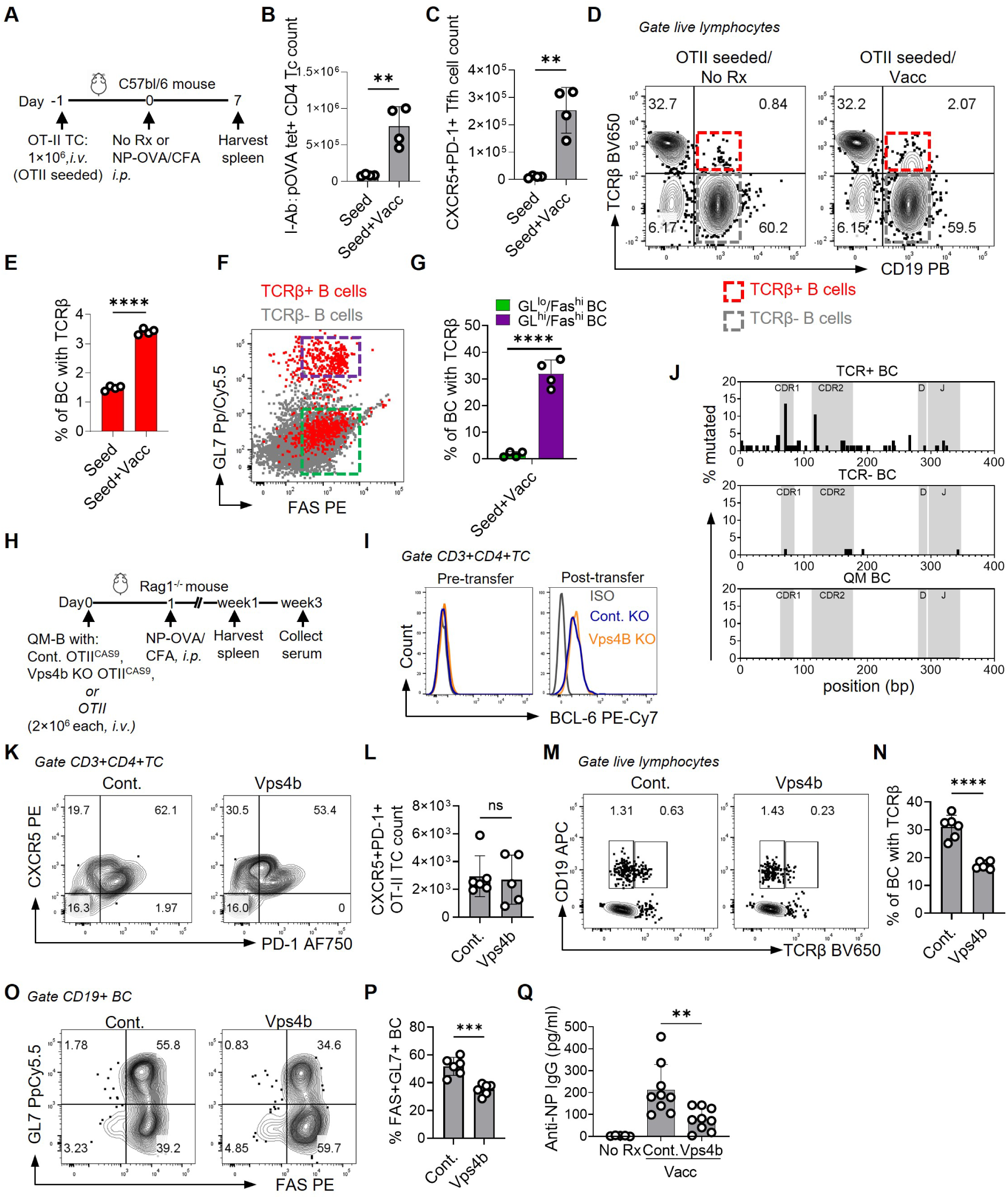
TCR transfer to B cells supports specific antibody production *in vivo*. (**A**) Schematic timeline of C57bl/6 immunization protocol to monitor T-B interactions and Tfh cell induction *in vivo*. To increase I-A^b^/pOVA-specific T cell precursor frequency, 10^6^ in vitro differentiated OT-II Th2 cells were adoptively transferred *i.v.* into mice 1 day before immunization. Mice were immunized *i.p.* with NP-OVA with CFA adjuvant, and splenocytes harvested one week post-vaccination for flow-cytometry analysis. (**B**), (**C**) Quantitation of I-A^b^/pOVA-specific CD4 T cells (B) and (C) Tfh cells, per spleen. Data points represent individual mice. Mean ± s.d is shown. Means were compared using two-tailed *t*-test. **(D)** Contour plots showing B cells with TCR transfer (TCRβ^+^ B cells, red dashed line and boxed gate) and B cells without TCR transfer (TCRβ^-^ BC, grey dashed and line boxed gate), from unvaccinated (No Rx) and vaccinated (Vacc) C57bl/6 mice, with seeded OT-II T cells (OTII seeded). (**E**) Quantitation of TCR transfer to B cells. Data points represent individual mice. Mean ± s.d is shown. Means were compared using two-tailed *t*-test. (**F**) Scatter plot overlay of Fas and GL7 expression on TCR^-^ and TCR^+^ B cells (gated as in (D)). OTII seeded mice left untreated (Seed) and after vaccination (Seed+Vacc). (**G**) Quantitation of TCR transfer to GL7^hi^ (purple dashed line and boxed region) and GL7^lo^ (green dashed line and boxed region) B cells. Data in (B), (C), (E), (G) are pooled from two independent experiments. (**H**) Schematic of Rag1^-/-^ adoptive transfer protocol timeline for T-B collaboration. Th2 polarized control or VPS4-deleted (VPS4 KO) OTII^CAS9^ T cell were transferred with QM-B cells. For V-region sequencing Th2 polarized OTII T cells were used. (**I**) Histogram overlays intracellular Bcl6 expression levels in the indicated OTII^CAS9^ Th2 cells before transfer into Rag^-/-^ mice, and 1 week post-inoculation with NP-OVA. (**J**) Plot of mutation frequency in genomic QM BCR heavy chain VDJ region, expressed as a percentage of total unique sequences detected by PCR. Shaded regions indicate complementarity determining regions (CDR). Splenic QM B cells with (TCR+ BC) and without (TCR-BC) transferred TCR were sorted 1 week post-inoculation with NP-OVA. Pre-transfer QM B cells (QM BC) were used as a baseline control. Results are pooled from two independent experiments. (**K**) Contour plots of CXCR5 and PD-1 expression levels in control (Cont.) or VPS4B-deleted (VPS4b) Th2 OTII^CAS9^ cells, isolated 1 week post-inoculation. (**L**) Quantitation of (K) expressed as cell numbers per spleen. (**M**) Contour plots labeling TCR (TCRβ) in CD19^+^ QM B cells cotransferred with the indicated OTII^CAS9^ Th2 cells. (**N**) Quantitation of (M). (**O**) Contour plots of Fas and GL7 expression levels in QM B cells cotransferred with the indicated OTII^CAS9^ Th2 cells. (**P**) Quantitation of (O). (**Q**) Quantitation of anti-NP IgG antibody levels in serum, as measured by ELISA, 3 weeks post-inoculation in untreated (No Rx) or in mice receiving cotransferred QM B cell and the indicated OTII^CAS9^ Th2 cells. For (L), (N), (P) and (Q), Data points represent individual mice. Mean ± s.d is shown. For (L), (N), (P) means were compared using two-tailed *t*-test. For (Q), means were compared by 1-way ANOVA corrected for multiple comparisons. ****, P < 0.0001; ***, P < 0.001; **, P < 0.01; ns, not significant (P > 0.05).

Due to technical limitations in isolating sufficient numbers of antigen-specific Tfh cells for *in vitro* CRISPR-mediated knockout of ECSRTs and isolation of Tfh cell-derived ectosome in bulk, we instead tested whether ectosomal TCR transfer to B cells affected antibody production *in vivo* using VPS4B-deleted *in vitro*-differentiated OTII^CAS9^ Th2 cells, in an adoptive transfer model using Rag^-/-^ mice as recipients (Fig. 4H). VPS4B-knockout or control OTII^CAS9^ Th2 cells were transferred intravenously into Rag^-/-^ mice, along with equal numbers of QM B cells, one day prior to immunization with NP-OVA (Fig. 4H). Surprisingly, transferred T cells 1 week post-inoculation exhibited an unexpected shift in phenotype, with up-regulation of Bcl-6 (Fig. 3I), and Tfh cell markers CXCR5 and PD-1 ((Fig. 4, K and L), revealing unexpected plasticity in their lineage commitment, and resulting in adaptive transdifferentiation *in vivo* to a Tfh-like phenotype (*44, 45*). This was not due to VPS4 deletion, as control T cells were equally affected, suggesting that cell-extrinsic mechanisms were responsible. TCR transfer to B cells was detectable in control OTII^CAS9^ T cells at the one-week timepoint, but was reduced by ∼50% in VPS4B-deleted T cells (Fig. 4, M and N). This was associated with the development of fewer GL7^+^Fas^+^ germinal center B cells (Fig. 4, O and P), indicating that germinal center responses were diminished without adequate ectosomal TCR transfer. Reduced TCR transfer to B cells resulted in a >50% reduction in NP-specific antibody levels measured in serum 3 weeks post-inoculation (Fig. 4Q), pointing to a substantial role for T cell ectosome-mediated help in B cell antibody responses *in vivo*.

## Synthetic reconstitution of ectosome-mediated help

We next sought to reconstitute ectosome-mediated help using minimal synthetic ectosomes (sEcto) only displaying recombinantly produced OTII TCR and CD40L (Fig. 5A). Single-chain constructs encoding OTII TCR V-domains (scTCR) (*46*), and mouse CD40L ectodomain fused to an oligomerization motif derived from the trimer-forming mutant of the GCN4 leucine zipper (*47, 48*), were expressed as fusion proteins with C-terminal SNAPf-tags (*49*) (Fig. 5,B and D; fig. S10, B and C). scTCR and CD40 fusion proteins were tethered to extruded DOPC liposomes, similar in size to native ectosomes (Fig. 5F), doped with benzylguanine- and fluorescent NBD-derivatized PE lipids (fig. S8, A and B).

**Fig. 5.**
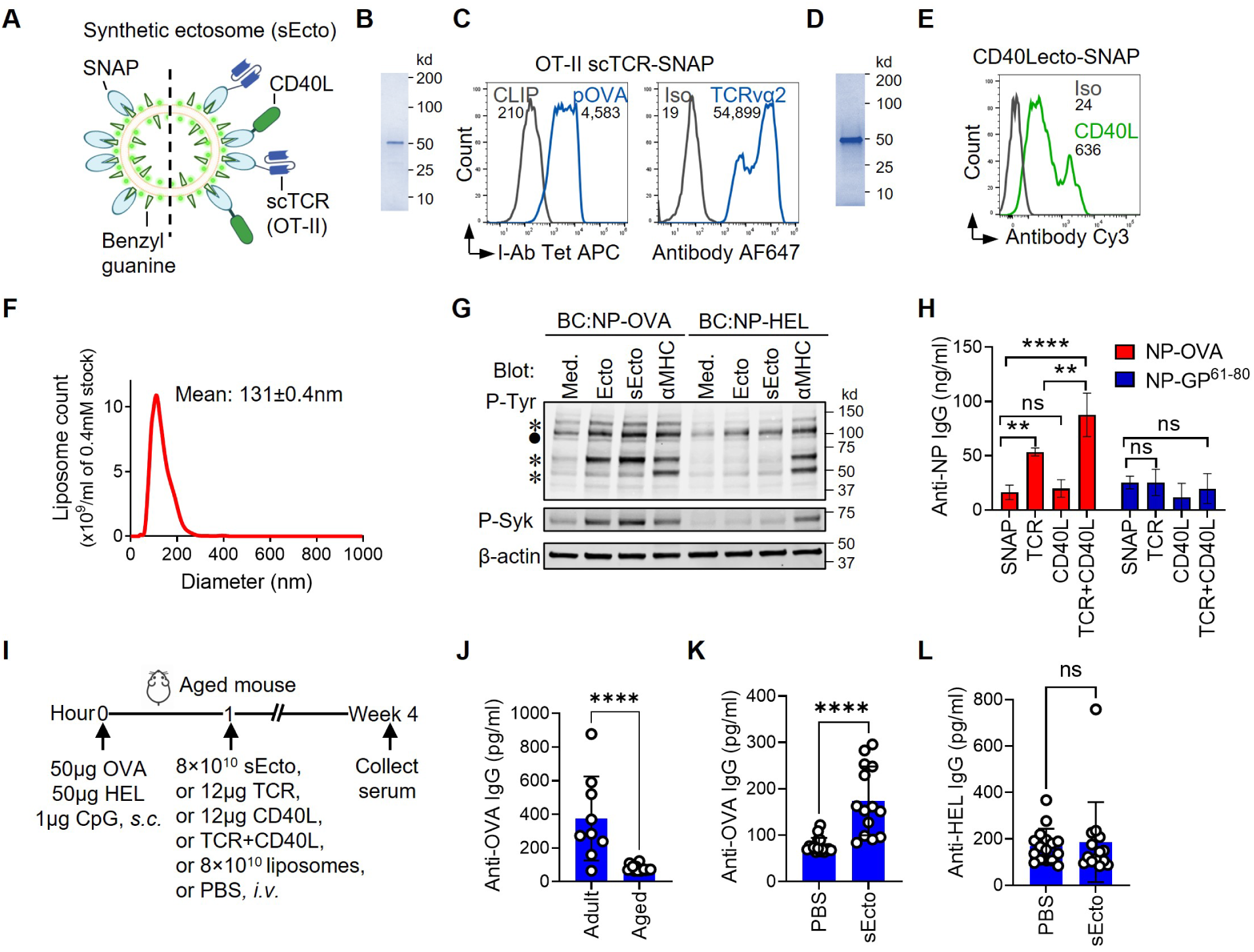
Biomimetic synthetic ectosomes reconstitute help for B cell antibody production. (**A**) Schematic of synthetic ectosomes made from extruded liposomes containing 10 mol% PE-NBD (light green dots), 10 mol% PE-benzylguanine (spikes) and 80 mol% DOPC lipids (base bilayer), to which single-chain OTII TCR and/or CD40L, recombinantly expressed as fusion proteins with C-terminal of SNAP-tags (SNAP), are covalently tethered (scOT-II-TCR-SNAP and CD40L-SNAP) (right of dashed line). SNAP-tags alone attached to liposomes were used as a negative control (left of dashed line). (**B**) Coomassie SDS-PAGE gel of purified OT-II-scTCR-SNAP. (**C**) Histogram overlays of specific binding of fluorescently labeled pOVA/I-A^b^ tetramer, or CLIP as a negative control, to detect OT-II-scTCR-SNAP attached to PE/DOPC bilayers formed on glass beads (left panel), or detection of OTII-scTCR-SNAP with a TCR Vα2-specific antibody (right panel). Numbers within panels represent mean fluorescence intensity. (**D**) Coomassie gel of purified CD40L-SNAP, and (**E**) Antibody labeling of bead-bilayer-attached CD40L-SNAP. (**F**) Size analysis of synthetic ectosomes by NTA. (**G**) Phosphotyrosine and pSyk Tyr352 immunoblot of lysates of NP-OVA-activated QM B cells (BC:NP-OVA), NP-HEL-activated QM B cells (BC:NP-HEL), or unstimulated QM B cells (Med.), incubated with OT-II T cell-derived ectosomes (Ecto), synthetic ectosomes (sEcto), or anti-MHC II-antibody (αMHC). Results are representative of 4 independent experiments. Asterisks indicate bands induced in NP-OVA activated B cells, filled circle represents band induced by ectosomes in both NP-OVA and NP-HEL-activated B cells. (**H**) IgG release by polyclonal memory-enriched B cells isolated from NP-BSA hyperimmunized C57bl/6 mice and pulsed with NP-OVA, or NP-GP^61-80^ as a non-specific control antigen, and cocultured with synthetic ectosomes displaying OT-II-scTCR-SNAP (TCR), mCD40L-SNAP or both. Liposomes displaying SNAP-tag alone were used as a negative control. Results are representative of 5 independent experiments. (**I**) Schematic of vaccination protocol timeline of aged mice vaccinated with OVA and HEL antigens with CpG adjuvant *s.c*., with or without subsequent *i.v.* infusion of synthetic ectosomes decorated with OT-II-scTCR-SNAP and CD40L-SNAP fusion proteins. (**J**) Quantitation of serum anti-OVA IgG in adult and aged mice post-vaccination; data are pooled from 3 independent experiments. Quantitation of serum IgG anti- OVA antibodies (**K**) and serum anti-HEL IgG antibodies (**L**) post-vaccination, with (sEcto) or without (PBS) synthetic ectosomes; data are pooled from 3 independent experiments. For J-L, data points represent individual animals. Mean ± s.d is shown. Means were compared using two-tailed *t*-test. ****, P < 0.0001; **, P < 0.01; *, P < 0.05; ns, not significant (P > 0.05).

Tethered proteins were detected on lipid bilayers, formed by liposome deposition on glass beads using flow-cytometry (Fig. 5, C and E) (*50*), or directly detected by single-particle flow-cytometry (fig S8, A to C). Fluorescently-tagged pOVA/I-A^b^ tetramers and conformation-sensitive anti-TCR Vα2 antibodies effectively labeled bilayer-tethered scOTII TCRs (Fig. 5C), indicating proper folding and intact pMHCII binding sites. To determine whether scOTII TCRs were functionally active, we attached them, along with His-tagged LFA-1 ligand ICAM-1, to glass supported planar lipid bilayers (SLB) containing PE-benzylguanine and DGS-NTA.Ni^3+^, to reconstitute a minimal binding surface for B cells (*51*) (Fig. S9A), and imaged B cells interacting with SLBs by TIRF microscopy (*52*). NP-OVA- but not NP-HEL-activated QM B cells triggered robust intracellular Ca^2+^-influx upon contact with scTCR-containing SLBs, while no response was detected on bilayers displaying SNAP-tag alone (fig. S9B), demonstrating that scTCRs could induced antigen-specific intracellular signaling in B cells. Similarly, bilayers displaying scTCR but not SNAP-tag accumulated I-A^b^ and induced protein tyrosine phosphorylation at the B cell contact interface (fig. S9, C to F).

Synthetic ectosomes displaying TCR and CD40L induced comparable patterns of protein tyrosine phosphorylation to OTII T cell-derived native ectosomes in QM B cells primed with IL- 4 and NP-OVA (Fig. 5G), including phosphorylation of Tyr352 in the activatory loop of Syk kinase (Fig. 5G). A similar pattern of protein tyrosine phosphorylation was induced by antibody-mediated MHCII crosslinking (Fig. 5G). In contrast, both native and synthetic ectosomes induced minimal protein phosphorylation in NP-HEL-primed B cells, demonstrating predominantly antigen-specific and TCR-dependent signaling in response to ectosomes. Taken together, these results point to TCR-mediated engagement and clustering of cognate pMHCII as the primary mechanism for physiological ectosome-evoked signal transduction.

To test the relative contributions of MHCII- and CD40-mediated signalling to B cell functional responses, we measured specific antibody secretion by MEBCs in response to synthetic ectosomes displaying scTCR, CD40L or both (fig. S8C). We also tethered the SNAP-tag polypeptide alone to liposomes as a non-binding control. CD40L-bearing synthetic ectosomes induced weak IgG antibody production by NP-OVA-primed MEBC, comparable to non-specific SNAP-tethered liposomes (Fig. 5H), suggesting that engagement of CD40L alone was insufficient for antibody production. A significant increase in IgG antibody production was detected in response to ectosomes displaying scTCR alone. However, ectosomes displaying both TCR and CD40 induced the highest levels of antibody release (Fig. 5H), suggesting synergistic crosstalk between pMHCII- and CD40-mediated signaling in B cell antibody responses. Antibody production by MEBC was highly antigen-specific, as only B cells primed with NP-OVA, and not NP-GFP-GP^61-80^, responded to scOTII TCR displaying ectosomes (Fig. 5H).

We next determined whether synthetic ectosomes augment T cell help to boost antibody responses to primary immunization in aged mice (Fig. 5I), which have severely compromised antibody responses to vaccination relative to healthy adult mice, mainly due to defective T helper function (*53, 54*) (Fig. 5J). We immunized aged C57bl/6 mice simultaneously with OVA and HEL, and infused synthetic ectosomes displaying both OTII scTCR and CD40L one hour after immunization, and measured serum antibody levels 4 weeks post-immunization (Fig. 5I). Addition of synthetic ectosomes resulted in highly significant enhancement of antibody production, increasing specific IgG levels by ∼2-fold (Fig. 5K), while no increase in anti-HEL antibodies was observed (Fig. 5L). Notably, administration of liposomes alone or soluble scTCR and CD40L in quantities comparable to the total protein or lipid dose of synthetic ectosomes failed to boost antibody production in coculture with NP-OVA-activated QM B cell (fig. S8D), or after immunization in aged mice (fig. S8E). These results demonstrate that intravenously administered synthetic ectosomes can penetrate SLO, and provide help to antigen-specific B cells through pMHCII and CD40 engagement, but only when TCR and CD40L are tethered to liposomes, suggesting that their attachment to lipid bilayers, which confers multivalency, lateral mobility, and binding-induced clustering, is critical for their function.

## Discussion

Our results demonstrate that differentiated T helper cells are superior to unpolarized CD4^+^ effector T cells in releasing ectosomes when activated *via* the TCR. These ectosomes are enriched in TCRs and key adhesion receptors, including SLAM family members and integrins, which are known to promote interactions with B cells, and are important for antibody responses (*28–31*). TCR-enriched ectosomes are transferred to antigen-activated B cells, where ectosome-tethered TCRs engage and cluster cognate pMHCII complexes, triggering intracellular signaling and class-switched antibody production. TCR engagement of pMHCII is generally understood to induce signaling primarily in T cells, while the nature and physiological relevance of pMHCII-mediated signals remains unresolved. Our results establish that engagement of cognate pMHCII by ectosome-tethered TCR induces B cell signaling and antibody production – a new modality of help for B cells that is distinct from direct T-B cell contact-dependent and soluble cytokine signals. We show, by deletion of the AAA+-ATPase VPS4B, which catalyses the terminal scission and release steps of nascent ectosomes from the plasma membrane (*32*), that ectosomes released from T helper cells account for ∼50% of the help delivered to B cells for antibody-production *in vitro*, and in an adoptive transfer mouse model of T-B cell collaboration. This implies that ectosomes released by T helper cells deliver signals for B cell help that are separable from those delivered through direct T-B cell interactions, since arrest of vesicle release only permits direct contact-dependent help, without ectosome-mediated contributions.

Our results suggest that T cell ectosomes that are released and transferred to B cells likely continue to deliver help after direct T-B interactions have terminated, prolonging signals beyond the often brief T-B cell encounters observed *in vivo* (fig. S11) (*55*). Consistent with this, we find that T cell ectosomes, together with helper cytokines, can support antibody production by B cells in the absence of T cells. While contact-dependent trans-synaptic transfer of ectosomes is likely to be most efficient (*23, 56*), released ectosomes may also escape the synaptic cleft and diffuse within SLO to provide help to bystander antigen-specific B cells (fig. S11). Moreover, the appearance of ectosome-like vesicles in the circulation of immunized mice raises the possibility that ectosomes mediate even longer range intercellular communication, transported in the circulation to anatomically distant tissues (fig. S11), protected from phagocytic clearance in transit by abundant CD47 on their surfaces (*57*) (Fig. 1E). Their presence in the circulation also makes them attractive candidates as readily accessible biomarkers of T cell help within less easily interrogable SLO. Our demonstration that ectosome-mediated help can be reconstituted by engineered synthetic ectosomes serves as a starting point for their development as a biomimetic nanoscale platform for therapeutic boosting of antibody responses, especially in conjunction with vaccines in the context of advanced age or CD4^+^ T cell immunodeficiency. Finally, other CD4^+^ T cell subsets which recognize pMHCII antigens, such as Tfr (*58*), Treg (*59*), and Th17 cells (*60*), likely also produce ectosomes with distinct effector compositions and cellular targets. Their detailed characterization will help in decoding the broader principles of ectosome-mediated intercellular communication with pMHCII-displaying antigen presenting cells, including B cells, monocyte/macrophages, dendritic cells (*24*) and activated human αβ (*61, 62*) and γδ T cells (*63*).

## Supporting information

Segmented tomogram of T cell ectosomes

## Acknowledgments

We thank Dr. Marilia Cascalho (U. Michigan) for the gift of NP-specific QM mice. I-A^b^/pOVA tetramers were obtained through the NIH Tetramer Core Facility. The support and resources from the University of Utah Arnold and Mabel Beckman Center for Cryo-EM and Center for High Performance Computing for cryo-EM and computational support, respectively, are gratefully acknowledged.

## Author contributions

K.C. conceived of the study, K.C. and F.L. designed the study, F.L. performed experiments, B.S., J.B., and W.I.S performed and analyzed cryo-EM and ECT experiments, F.L. and H.R. performed proteomics experiments and H.R. advised on analysis, all authors contributed to editing the final manuscript.

## Data and materials availability

All data associated with the present study are available from the corresponding author upon request.

## Supplementary Materials

Materials and Methods

Figs. S1 to S11

Tables S1 and S2

Movies S1

## References

1. D. DiToro, N. Murakami, S. Pillai, T-B Collaboration in Autoimmunity, Infection, and Transplantation. Transplantation 108, 386–398 (2024).

2. A. Risso, M. E. Cosulich, A. Rubartelli, M. R. Mazza, A. Bargellesi, MLR3 molecule is an activation antigen shared by human B, T lymphocytes and T cell precursors. Eur J Immunol 19, 323–328 (1989).

3. P. A. Roche, K. Furuta, The ins and outs of MHC class II-mediated antigen processing and presentation. Nat Rev Immunol 15, 203–216 (2015).

4. D. J. Lenschow et al., Differential up-regulation of the B7-1 and B7-2 costimulatory molecules after Ig receptor engagement by antigen. J Immunol 153, 1990–1997 (1994).

5. C. Dong, Cytokine Regulation and Function in T Cells. Annu Rev Immunol 39, 51–76 (2021).

6. D. van Essen, H. Kikutani, D. Gray, CD40 ligand-transduced co-stimulation of T cells in the development of helper function. Nature 378, 620–623 (1995).

7. W. J. Poo, L. Conrad, C. A. Janeway, Jr., Receptor-directed focusing of lymphokine release by helper T cells. Nature 332, 378–380 (1988).

8. A. Grakoui et al., The immunological synapse: a molecular machine controlling T cell activation. Science 285, 221–227 (1999).

9. J. C. Cambier et al., Ia binding ligands and cAMP stimulate nuclear translocation of PKC in B lymphocytes. Nature 327, 629–632 (1987).

10. N. Nabavi et al., Signalling through the MHC class II cytoplasmic domain is required for antigen presentation and induces B7 expression. Nature 360, 266–268 (1992).

11. P. J. Lane, F. M. McConnell, G. L. Schieven, E. A. Clark, J. A. Ledbetter, The role of class II molecules in human B cell activation. Association with phosphatidyl inositol turnover, protein tyrosine phosphorylation, and proliferation. J Immunol 144, 3684–3692 (1990).

12. P. Lang et al., TCR-induced transmembrane signaling by peptide/MHC class II via associated Ig-alpha/beta dimers. Science 291, 1537–1540 (2001).

13. J. C. Cambier, K. R. Lehmann, Ia-mediated signal transduction leads to proliferation of primed B lymphocytes. J Exp Med 170, 877–886 (1989).

14. G. A. Bishop, W. D. Warren, M. T. Berton, Signaling via major histocompatibility complex class II molecules and antigen receptors enhances the B cell response to gp39/CD40 ligand. Eur J Immunol 25, 1230–1238 (1995).

15. R. Guy et al., Antigen-specific helper function of cell-free T cell products bearing TCR V beta 8 determinants. Science 244, 1477–1480 (1989).

16. R. Guy, R. J. Hodes, Antigen-specific, MHC-restricted B cell activation by cell-free Th2 cell products. Synergy between antigen-specific helper factors and IL-4. J Immunol 143, 1433–1440 (1989).

17. K. Choudhuri et al., Polarized release of T-cell-receptor-enriched microvesicles at the immunological synapse. Nature 507, 118–123 (2014).

18. D. G. Saliba et al., Composition and structure of synaptic ectosomes exporting antigen receptor linked to functional CD40 ligand from helper T cells. Elife 8, (2019).

19. M. J. Barnden, J. Allison, W. R. Heath, F. R. Carbone, Defective TCR expression in transgenic mice constructed using cDNA-based alpha- and beta-chain genes under the control of heterologous regulatory elements. Immunol Cell Biol 76, 34–40 (1998).

20. M. Cascalho, A. Ma, S. Lee, L. Masat, M. Wabl, A quasi-monoclonal mouse. Science 272, 1649–1652 (1996).

21. C. Thery et al., Minimal information for studies of extracellular vesicles 2018 (MISEV2018): a position statement of the International Society for Extracellular Vesicles and update of the MISEV2014 guidelines. J Extracell Vesicles 7, 1535750 (2018).

22. F. G. Kugeratski et al., Quantitative proteomics identifies the core proteome of exosomes with syntenin-1 as the highest abundant protein and a putative universal biomarker. Nat Cell Biol 23, 631–641 (2021).

23. J. C. Stinchcombe et al., Ectocytosis renders T cell receptor signaling self-limiting at the immune synapse. Science 380, 818–823 (2023).

24. P. F. Cespedes et al., T-cell trans-synaptic vesicles are distinct and carry greater effector content than constitutive extracellular vesicles. Nat Commun 13, 3460 (2022).

25. R. D. Fisher et al., Structural and biochemical studies of ALIX/AIP1 and its role in retrovirus budding. Cell 128, 841–852 (2007).

26. O. Schmidt, D. Teis, The ESCRT machinery. Curr Biol 22, R116–120 (2012).

27. C. Doyle, J. L. Strominger, Interaction between CD4 and class II MHC molecules mediates cell adhesion. Nature 330, 256–259 (1987).

28. H. Qi, J. L. Cannons, F. Klauschen, P. L. Schwartzberg, R. N. Germain, SAP-controlled T-B cell interactions underlie germinal centre formation. Nature 455, 764–769 (2008).

29. J. L. Cannons et al., Optimal germinal center responses require a multistage T cell:B cell adhesion process involving integrins, SLAM-associated protein, and CD84. Immunity 32, 253–265 (2010).

30. M. C. Zhong et al., SLAM family receptors control pro-survival effectors in germinal center B cells to promote humoral immunity. J Exp Med 218, (2021).

31. C. R. Monks, B. A. Freiberg, H. Kupfer, N. Sciaky, A. Kupfer, Three-dimensional segregation of supramolecular activation clusters in T cells. Nature 395, 82–86 (1998).

32. J. E. Garrus et al., Tsg101 and the vacuolar protein sorting pathway are essential for HIV- 1 budding. Cell 107, 55–65 (2001).

33. R. J. Platt et al., CRISPR-Cas9 knockin mice for genome editing and cancer modeling. Cell 159, 440–455 (2014).

34. S. B. Gauld, J. M. Dal Porto, J. C. Cambier, B cell antigen receptor signaling: roles in cell development and disease. Science 296, 1641–1642 (2002).

35. G. Grynkiewicz, M. Poenie, R. Y. Tsien, A new generation of Ca2+ indicators with greatly improved fluorescence properties. J Biol Chem 260, 3440–3450 (1985).

36. N. M. Weisel et al., Surface phenotypes of naive and memory B cells in mouse and human tissues. Nat Immunol 23, 135–145 (2022).

37. A. Oxenius, M. F. Bachmann, R. M. Zinkernagel, H. Hengartner, Virus-specific MHC-class II-restricted TCR-transgenic mice: effects on humoral and cellular immune responses after viral infection. Eur J Immunol 28, 390–400 (1998).

38. G. D. Victora, M. C. Nussenzweig, Germinal Centers. Annu Rev Immunol 40, 413–442 (2022).

39. S. Crotty, Follicular helper CD4 T cells (TFH). Annu Rev Immunol 29, 621–663 (2011).

40. S. Crotty, T Follicular Helper Cell Biology: A Decade of Discovery and Diseases.Immunity 50, 1132–1148 (2019).

41. K. G. Smith, G. J. Nossal, D. M. Tarlinton, FAS is highly expressed in the germinal center but is not required for regulation of the B-cell response to antigen. Proc Natl Acad Sci U S A 92, 11628–11632 (1995).

42. L. Cervenak, A. Magyar, R. Boja, G. Laszlo, Differential expression of GL7 activation antigen on bone marrow B cell subpopulations and peripheral B cells. Immunol Lett 78, 89–96 (2001).

43. J. M. Di Noia, M. S. Neuberger, Molecular mechanisms of antibody somatic hypermutation. Annu Rev Biochem 76, 1–22 (2007).

44. A. Glatman Zaretsky et al., T follicular helper cells differentiate from Th2 cells in response to helminth antigens. J Exp Med 206, 991–999 (2009).

45. W. Song, J. Craft, T Follicular Helper Cell Heterogeneity. Annu Rev Immunol 42, 127–152 (2024).

46. K. S. Gunnarsen et al., Periplasmic expression of soluble single chain T cell receptors is rescued by the chaperone FkpA. BMC Biotechnol 10, 8 (2010).

47. P. B. Harbury, T. Zhang, P. S. Kim, T. Alber, A switch between two-, three-, and four-stranded coiled coils in GCN4 leucine zipper mutants. Science 262, 1401–1407 (1993).

48. N. Levin et al., Spontaneous Activation of Antigen-presenting Cells by Genes Encoding Truncated Homo-Oligomerizing Derivatives of CD40. J Immunother 40, 39–50 (2017).

49. X. Sun et al., Development of SNAP-tag fluorogenic probes for wash-free fluorescence imaging. Chembiochem 12, 2217–2226 (2011).

50. T. J. Crites et al., Supported Lipid Bilayer Technology for the Study of Cellular Interfaces. Curr Protoc Cell Biol 68, 24 25 21–24 25 31 (2015).

51. D. Depoil et al., CD19 is essential for B cell activation by promoting B cell receptor-antigen microcluster formation in response to membrane-bound ligand. Nat Immunol 9, 63–72 (2008).

52. C. Beilin et al., Dendritic cell-expressed common gamma-chain recruits IL-15 for trans-presentation at the murine immunological synapse. Wellcome Open Res 3, 84 (2018).

53. P. T. Sage, C. L. Tan, G. J. Freeman, M. Haigis, A. H. Sharpe, Defective TFH Cell Function and Increased TFR Cells Contribute to Defective Antibody Production in Aging. Cell Rep 12, 163–171 (2015).

54. A. Silva-Cayetano et al., Spatial dysregulation of T follicular helper cells impairs vaccine responses in aging. Nat Immunol 24, 1124–1137 (2023).

55. Z. Shulman et al., Dynamic signaling by T follicular helper cells during germinal center B cell selection. Science 345, 1058–1062 (2014).

56. T. Igakura et al., Spread of HTLV-I between lymphocytes by virus-induced polarization of the cytoskeleton. Science 299, 1713–1716 (2003).

57. P. A. Oldenborg et al., Role of CD47 as a marker of self on red blood cells. Science 288, 2051–2054 (2000).

58. M. A. Linterman et al., Foxp3+ follicular regulatory T cells control the germinal center response. Nat Med 17, 975–982 (2011).

59. I. S. Okoye et al., MicroRNA-Containing T-Regulatory-Cell-Derived Exosomes Suppress Pathogenic T Helper 1 Cells. Immunity 41, 503 (2014).

60. Ivanov, II et al., The orphan nuclear receptor RORgammat directs the differentiation program of proinflammatory IL-17+ T helper cells. Cell 126, 1121–1133 (2006).

61. A. Lanzavecchia, E. Roosnek, T. Gregory, P. Berman, S. Abrignani, T cells can present antigens such as HIV gp120 targeted to their own surface molecules. Nature 334, 530–532 (1988).

62. H. S. Ko, S. M. Fu, R. J. Winchester, D. T. Yu, H. G. Kunkel, Ia determinants on stimulated human T lymphocytes. Occurrence on mitogen- and antigen-activated T cells. J Exp Med 150, 246–255 (1979).

63. M. Brandes, K. Willimann, B. Moser, Professional antigen-presentation function by human gammadelta T Cells. Science 309, 264–268 (2005).

64. A. Keppler et al., A general method for the covalent labeling of fusion proteins with small molecules in vivo. Nat Biotechnol 21, 86–89 (2003).

65. J. Shine, L. Dalgarno, Occurrence of heat-dissociable ribosomal RNA in insects: the presence of three polynucleotide chains in 26 S RNA from cultured Aedes aegypti cells. J Mol Biol 75, 57–72 (1973).

66. M. Naito et al., CD40L-Tri, a novel formulation of recombinant human CD40L that effectively activates B cells. Cancer Immunol Immunother 62, 347–357 (2013).

67. A. Oxenius et al., Presentation of endogenous viral proteins in association with major histocompatibility complex class II: on the role of intracellular compartmentalization, invariant chain and the TAP transporter system. Eur J Immunol 25, 3402–3411 (1995).

68. E. Dekel et al., Antibody-Free Labeling of Malaria-Derived Extracellular Vesicles Using Flow Cytometry. Biomedicines 8, (2020).

69. D. N. Mastronarde, Automated electron microscope tomography using robust prediction of specimen movements. J Struct Biol 152, 36–51 (2005).

70. G. C. Lander et al., Appion: an integrated, database-driven pipeline to facilitate EM image processing. J Struct Biol 166, 95–102 (2009).

71. A. J. Noble, S. M. Stagg, Automated batch fiducial-less tilt-series alignment in Appion using Protomo. J Struct Biol 192, 270–278 (2015).

72. H. Winkler, K. A. Taylor, Accurate marker-free alignment with simultaneous geometry determination and reconstruction of tilt series in electron tomography. Ultramicroscopy 106, 240–254 (2006).

73. J. I. Agulleiro, J. J. Fernandez, Fast tomographic reconstruction on multicore computers. Bioinformatics 27, 582–583 (2011).

74. J. I. Agulleiro, J. J. Fernandez, Tomo3D 2.0--exploitation of advanced vector extensions (AVX) for 3D reconstruction. J Struct Biol 189, 147–152 (2015).

75. T. Grant, N. Grigorieff, Measuring the optimal exposure for single particle cryo-EM using a 2.6 A reconstruction of rotavirus VP6. Elife 4, e06980 (2015).

76. J. R. Kremer, D. N. Mastronarde, J. R. McIntosh, Computer visualization of three-dimensional image data using IMOD. J Struct Biol 116, 71–76 (1996).

77. Y. T. Liu et al., Isotropic reconstruction for electron tomography with deep learning. Nat Commun 13, 6482 (2022).

78. E. F. Pettersen et al., UCSF ChimeraX: Structure visualization for researchers, educators, and developers. Protein Sci 30, 70–82 (2021).

79. J. Schindelin et al., Fiji: an open-source platform for biological-image analysis. Nat Methods 9, 676-682 (2012).

80. K. Choudhuri, D. Wiseman, M. H. Brown, K. Gould, P. A. van der Merwe, T-cell receptor triggering is critically dependent on the dimensions of its peptide-MHC ligand. Nature 436, 578–582 (2005).

81. B. Zybailov et al., Statistical analysis of membrane proteome expression changes in Saccharomyces cerevisiae. J Proteome Res 5, 2339–2347 (2006).

82. K. Tzelepis et al., A CRISPR Dropout Screen Identifies Genetic Vulnerabilities and Therapeutic Targets in Acute Myeloid Leukemia. Cell Rep 17, 1193–1205 (2016).

83. K. Labun et al., CHOPCHOP v3: expanding the CRISPR web toolbox beyond genome editing. Nucleic Acids Res 47, W171–W174 (2019).

84. K. M. Toellner et al., Low-level hypermutation in T cell-independent germinal centers compared with high mutation rates associated with T cell-dependent germinal centers. J Exp Med 195, 383–389 (2002).

85. M. L. Dustin, T. Starr, R. Varma, V. K. Thomas, Supported planar bilayers for study of the immunological synapse. Curr Protoc Immunol **Chapter** 18, 18 13 11–18 13 35 (2007).

86. R. Varma, G. Campi, T. Yokosuka, T. Saito, M. L. Dustin, T cell receptor-proximal signals are sustained in peripheral microclusters and terminated in the central supramolecular activation cluster. Immunity 25, 117–127 (2006).

