## Supplemental Information for "T cell ectosomes promote antibody responses through cognate TCR-pMHC interactions"

**The PDF file includes:**

Materials and Methods
Figs. S1 to S11
Tables S1 and S2
References

**Other Supplementary Materials for this manuscript include the following:**

Movie S1

### Materials and Methods

#### Expression constructs and recombinant proteins

OT-II TCR was expressed as single-chain variable domain fused to a sequence encoding the SNAP-tag polypeptide (*64*) in *E.coli* using a method adapted from Gunnarsen *et al* (*46*). The pelB signal peptide was at the N-terminus of expression constructs to promote periplasmic translocation and disulphide bond formation. This was followed sequentially by segments coding for the OT-II Vα segment, a synthetic 21 amino acid linker (KLSGSASAPKLEEGEFSEARV (*46*)), the corresponding TCR Vβ segment, a cMyc-tag for immunodetection, a flexible (G4S)2 linker, a SNAP-tag sequence, a TEV protease cleavage site, and a C-terminal 12×His-tag. To increase scTCR expression levels, a periplasmic chaperon FkpA was co-expressed by introducing a Shine–Dalgarno (SD) sequence (*65*) between the scTCR-SNAP-tag frame and FkpA frame. Coding sequences for the all segments were synthesized as a codon-optimized gBlock DNA duplexes and inserted into XbaI/NotI linearized pET vector after the T7 promoter by In-Fusion cloning (Clontech). For scTCR-SNAP protein expression, pET constructs were transfected into chemically competent *E.coli* (BL21 Star™ (DE3) pLysS One Shot™, Invitrogen), and cultures expanded to $OD^{600}$ ~ 0.5 at 37°C, prior to induction with IPTG (0.1mM for OT-II TCR) for 16 hours at 18°C. *E.coli* was harvested and resuspended in 25 mM Hepes, 150 mM KCl, and 10 mM imidazole, pH 7.4, and lysed using 0.1mg/ml lysozyme and 20μg/ml DNase, followed by sonication. After centrifugation at 10,000g for 30 minutes, culture supernatant was suction-filtered (0.45μm pore size) and affinity-purified by two rounds of imidazole gradient elution over a HiTrap TALON crude column (Cytiva) on a AKTA Pure FPLC system (GE Healthcare). C-terminal His-tags were removed by TEV protease digestion (1000U/mg protein) for 16 hours at 4°C, and poly-His fragments removed using a HiTrap TALON column. The flow-though was concentrated and subjected to a final polishing and buffer exchange step on a Superdex™ 75 Increase gel-filtration column (Cytiva) yielding scTCR-SNAP proteins of the expected molecular weight at >94% purity.

Mouse CD40L-SNAP fusion protein was cloned in-frame into the $pSNAP_f$ vector (NEB) (*49*). For protein secretion, the human IL-2 signal peptide was inserted after the CMV promoter, followed sequentially by coding sequences for a 12×His-tag, a TEV cleavage site, SNAP-tag, a Flag-tag for immunodetection, a trimeric leucine zipper GCN4 (*66*), (G2TG2S)3 linker, and the mouse CD40L ectodomain (aa50-260). Codon-optimized DNA gBlock duplexes encoding the various fragments were inserted into AgeI/NotI linearized $pSNAP_f$ vector by In-Fusion cloning. The ExpiCHO™ Expression System was used to express CD40L-SNAP protein following the "max titer" protocol suggested by the manufacturer (Gibco). Briefly, ExpiCHO-S cells were expanded to $1.2\times10^9$ cells in 200ml ExpiCHO™ Expression Medium in a 1L vented plastic erlenmeyer flask, before being transfected with 200μg $pSNAP_f$-CD40L plasmid and 640μL ExpiFectamine CHO reagent for 20 hours at 37°C, 5% $CO_2$. After transfection, cells were placed on an orbital shaker set at 1000rpm in a 32°C incubator, 5% $CO_2$. 1.2ml ExpiCHO Enhancer was added on day 1 of incubation, and 32ml ExpiCHO Feed was added on days 1 and 5. Supernatant was harvested on day 12-14, and buffer-exchanged by dialysis (30kD cut-off) into 25 mM Hepes, 150 mM KCl, and 10 mM imidazole, pH 7.4, prior to purification as described for TCR-SNAP proteins.

A fusion protein of the LCMV-$GP^{61-80}$ (I-$A^b$-restricted peptide epitope) (*67*) and EGFP was synthesised as codon optimized gBlock duplexes encoding $GP^{61-80}$ fused to EGFP and a 12×His-tag sequence, and cloned into XhoI/NotI digested pET vector. The *E.coli* transfection, protein induction and purification were similar to the scTCR, the only difference was that the induction was by 0.5mM IPTG for 5 hours at 37 °C. Protein purity was assayed via SDS-PAGE. Construct maps are shown in fig. S10.

**Mice**

C57bl/6J mice, OT-II transgenic mice (B6.Cg-Tg(TcraTcrb)425Cbn/J), Rag1$^{-/-}$ mice (B6.129S7-*Rag1*$^{tm1Mom}$/J), Rosa26-Cas9 knock-in mice (B6J.129(Cg)-*Gt(ROSA)26Sor*$^{tm1.1(CAG-cas9*,-EGFP)Fezh}$/J), CD45.1$^{+}$ C57bl/6J mice (B6.SJL-*Ptprc*$^{a}$ *Pepc*$^{b}$/BoyJ) and SMARTA-1 mice (B6.Cg-*Ptprc*$^{a}$ *Pepc*$^{b}$ Tg(TcrLCMV)1Aox/PpmJ) were purchased from Jackson Laboratory. QM transgenic mice were a gift from Dr. Marilia Cascalho (University of Michigan Medical School) (*20*). All mice were bred and maintained under specific pathogen-free (SPF) housing conditions at the University of Michigan Medical School animal housing facility, in accordance with Institutional Animal Care & Use Committee (IACUC)-approved protocols (PRO00010222). Mice were maintained and used in experiments in accordance with local, state, federal, and NIH regulations.

**Cell isolation and purification**

Spleens were harvested and rinsed with MACS buffer (2% FCS/PBS, 1mM EDTA) and gently macerated with the end of a 3 ml syringe. The resulting cell suspension was filtered through a 40µm cell strainer and pelleted by centrifugation. For flow-cytometry analysis and *in vitro* culture, splenocytes were treated with RBC lysis buffer for 5 minutes and filtered again to remove membrane debris. Otherwise, the cells were resuspended in MACS buffer for downstream negative isolation. Mouse CD4$^{+}$ T cells and B cells were negatively purified by EasySep™ Mouse CD4$^{+}$ T Cell Isolation Kit (Stemcell) and EasySep™ Mouse B Cell Isolation Kit (Stemcell), respectively.

**Th2 cell differentiation and ectosome production**

To generate Th2-differentiated OT-II cells, splenocytes from OT-II transgenic mice were cultured in RPMI-1640 medium containing 1µM OVA$^{323-339}$ peptide (Anaspec), 10% exosome-depleted FCS (Gibco), 100u/ml hIL-2, 10ng/ml mIL-4, 5µg/ml anti-IL-12 p40, 5µg/ml anti-IFNγ, 50µM 2ME and 100U/ml penicillin-streptomycin for 7 days. Live CD4$^{+}$ T cells were negatively enriched by MACS isolation (Stemcell) and rested in complete exosome-depleted medium containing 10U/ml IL-2 for 5-7 days until quiescent. Th2 Smarta-T cells were induced similarly, using 0.2µM LCMV GP$^{61-80}$ peptide (Anaspec). Th2 differentiation was confirmed by IL-4 release upon TCR stimulation, and/or by Gata-3 expression detected by flow-cytometry. Undifferentiated T cells (Th0) were used as a control and were cultured without polarizing factors IL-4, anti-IL-12 p40, or anti-IFNγ.

Quiescent Th2 or Th0 CD4$^{+}$ T cells were stimulated by plate-bound anti-CD3ε (145-2C11, 5µg/ml) with or without anti-CD28 (37.51, 5µg/ml) or recombinant pOVA/I-A$^{b}$ monomers (1.5µg/ml, NIH Tetramer Core Facility, Emory University) with or without mCD80 (5µg/ml, SinoBiological). Alternatively, T cells were activated by phorbol myristate acetate (PMA, 50ng/ml) and ionomycin (1µM), or anti-CD3ε (145-2C11, 10µg/ml), in solution at the indicated concentrations for 48 or 72 hours. For ectosome isolation, activated T cell culture supernatants were collected and subjected to serial centrifugation at 500g for 30 minutes, 2,000g for 30 minutes, and 10,000g for 30 minutes, to remove live cells, dead cells and other small cellular debris respectively. Ectosomes in culture supernatants were pelleted and resuspended in PBS by two rounds of ultracentrifugation at 100,000g for 120 minutes, and finally resuspended in approximately 50µl PBS or culture medium for downstream experiments. A workflow was shown in fig. S1A.

**Nanoparticle tracking analysis (NTA)**

Ectosomes or liposomes were diluted to approximately $10^8$ particles/ml in filter-degassed PBS before detection using a NanoSight NS300 instrument (Malvern Panalytical). The samples were analyzed using a 488nm laser (camera level 16) and videos were recorded at 25fps for 60 seconds, with 5 videos/sample. The analysis detection threshold was set to 3. The binned data or mean density was plotted in GraphPad Prism 9 for comparison between samples.

To deplete $TCR^+$ ectosomes, $1\times10^9$ Ectosomes produced by activated OT-II T cells were incubated with 0.2mg/ml of biotinylated anti-CD3ε antibody (17A2) or an isotype control antibody in 250µl of PBS for 1 hour at room temperature. Then, 50µl of Streptavidin-MojoBeads (Biolegend) were added for an additional 2 hour incubation. Magnet beads were removed from samples by exposure to a DynaMag™-Spin Magnet (Invitrogen), followed by centrifugation at 10,000g for 30 minutes. The resulting supernatant was collected for quantitation of ectosomes as described above.

### Small-particle flow-cytometry

To detect ectosome and associated fluorescently-labeled surface proteins, small-particle flow-cytometry was performed using a Ze5 flow cytometer (Bio-Rad), equipped with a secondary small particle forward-scatter detector along with a high power 405nm laser line to maximize scattering signal(*68*). Particle-associated fluorescence was detected using a 488nm laser line with 509/12nm emission filter, a 561nm line with 577/8nm emission filter, and a 640nm laser with 670/15nm emission filter. An inline 0.2µm filter was installed to reduce instrument and background noise. Buffers were filtered through a 100nm filter, degassed under vacuum overnight, and ultracentrifuged at 100,000g overnight to remove small-size impurities and bubbles. Tubes and pipette tips were rinsed with freshly degassed and ultracentrifuged buffer prior to use. Ectosomes were first stained with the lipid dye PKH67 (Sigma) following the manufacturer's instructions. Briefly, the exosome pellet was stained with 6µl of PKH67 in 2ml of "Diluent C" for 10 minutes. It was then quenched by adding 2ml of 10% BSA/PBS, diluted with 4.5ml of medium, and placed on top of 1.5ml of 0.971M sucrose, and ultracentrifuged at 192,000g for 2 hours. The pellet was washed through a 100kD Amicon centrifugal filter (Sigma) to remove any remaining traces of free PKH67 dye. The PKH67-stained ectosomes were resuspended in PBS and stained with 20µg/ml of fluorescently labeled antibodies (conjugated with AF647 and Cy3 in-house, F/P ratio >7) at room temperature for 1 hour. Samples were then washed by resuspension in PBS and ultracentrifugation at 192,000g for 1 hour. The resulting pellet was resuspended in 500µL of PBS. After extensive washing the Ze5 with degassed and ultracentrifuged PBS, samples were acquired at low speed (0.1µl/sec) to avoid the turbulence-induced particle "swarming". The flow cytometry files were analyzed using Flowjo10, and 99% of events above the instrument noise were $PKH67^+$. A similar protocol was used for analysis of both OT-II-derived ectosomes (ECTO) and synthetic T cell-mimetic ectosomes (sEcto).

### Plasma ectosome isolation

8-10-week-old C57BL/6 mice were inoculated *s.c.* with 100 µg of OVA/CFA for the first round of immunization, followed by 100 µg OVA/IFA for subsequent boosts, at two-week intervals. One week after each vaccination, blood samples were collected into EDTA-coated tubes. After gently mixing, the samples were left at RT for 15mins and then subjected to serial centrifugations (500g×10min, 2,000g×10min, and 2,000g×10min) to remove live/dead cells. After filtering through a 0.2µm filter, the plasma was then subjected to centrifugation at 10,000g for 10 minutes to remove protein debris and large particles. Finally, the remaining ectosomes were pelleted using an ultracentrifuge at 100,000g for 2hrs, and resuspended in PBS for subsequent downstream analysis as described in ***Small-particle flow-cytometry*** section.

**Transmission cryoelectron microscopy (cryo-EM) and electron cryotomography (ECT) of OTII vesicles**

OTII-derived vesicles were prepared for cryo-EM and ECT by incubating 3.5 µl of a concentrated vesicle suspension (200µl PBS containing ECTOSOMES derived from 40 million OTII T cells stimulated by plate-bound αCD3ε/CD28 or PBS) on a freshly glow-discharged (PELCO easiGlow glow discharge system, 25 mA, 10 seconds) 300-mesh Lacey Carbon EM grid (PELCO NetMesh). Samples were incubated for one minute within the environmental chamber of a Leica GP2 plunge freezer (4 °C, 95% relative humidity), excess liquid was removed by blotting (Ted Pella 595 filter paper, 3.5 seconds), and samples were plunge frozen in liquid ethane.

Tilt-series were acquired on a Titan Krios (300 keV) equipped with a Gatan K3 Summit direct electron detector and a post-column Gatan Bioquantum energy filter. Tilt series were collected in super-resolution mode using SerialEM (*69*) (bidirectionally starting from 18° moving to -60° and then 21° to 60° in 3° step sizes, 188 ms total exposure for each tilt image) at a nominal magnification of 81,000×, corresponding to a pixel size of 0.53 Å, with defocus ranging between 2 and 6 microns, and a total dose per specimen 121 $e^{-}/Å^{2}$. Full-frame alignments were performed using SerialEM without dose weighting.

Tilt-series were aligned using Appion-Protomo (*70-72*) after downsampling by a binning factor of two, corresponding to a pixel size of 1.06Å. After initial coarse alignment, each tilt-series was manually aligned, and then refined using a set of alignment thicknesses between 800 and 1,500Å. Automated and manual refinements were iterated until the refinements converged, and then reconstructed for visual analysis using Tomo3D SIRT (*73, 74*) after moderate dose-compensation using the relation described in (*75*). CTF correction was not performed at this stage. 3dmod (*76*) was used to prepare the tomogram slices (e.g., as shown in Fig. 1d).

Isosurface rendering and missing-wedge corrections were performed using Isonet (*77*). Briefly, reconstructed tomograms were downsampled using a binning factor of 16 and the CTFs were deconvolved using Isonet. CTF-corrected tomograms were imported as a training set into Isonet's deep-learning neural network and used to iteratively reconstruct the missing-wedge information (*77*). The corrected tomograms were then scaled to match the pixel size of the original tomograms. Isosurface renderings, generated by Isonet, were modelled onto the original tomograms, using ChimeraX (*78*), and used to generate the image shown in Fig. 1d, and in the Supplementary Movie.

Numbers of vesicles released from antigen-stimulated and unstimulated OTII cells were quantified from projection cryo-EM images collected at a nominal magnification of 6100× (to a pixel size of 41.17Å). Four randomly selected projection images were visualized in FIJI (*79*), and the number of vesicles per image was counted manually to generate the data shown in fig. S2I. Mean values and standard deviations between the four representative fields chosen at random were calculated, and means were compared for significance using two-tailed *t*-test, using GraphPad Prism. Vesicle diameter was measured from a series of projection images collected in super-resolution mode (nominal magnification of 81,000×, corresponding to a pixel size of 0.53Å, defocus range of 1.5 - 2.5 microns). Images were visualized in FIJI, and diameters were measured using the FIJI measure tool. 50 vesicles were chosen at random and measured for each sample to generate the data in fig. S2J, mean values and standard deviations were calculated, and means compared for significance using two-tailed *t*-test, using GraphPad Prism.

**Western blot and biochemistry**

Immunoblotting was performed as previously described (*80*). Briefly, T cells, ectosomes or B cells stimulated with the indicated reagents were lysed using chilled RIPA buffer (1% Triton X-100, 140mM NaCl, 50mM Hepes, 10mM Idoacetamide, 1×MS-SAFE Protease and Phosphatase Inhibitor (Sigma), pH 7.4). The reduced lysate was separated by SDS-PAGE, transferred to nitrocellulose membranes, and probed with specific antibodies (listed in Table S1) for the signaling and (loading) control proteins. The blots were visualized using the Odyssey® Classic Infrared Imaging System and processed using Fiji/ImageJ 1.53. Some blots were stripped using NewBlot™ stripping buffer (Licor) and reprobed.

**Proteomic analysis of ectosomes**

Proteomic analysis was performed in collaboration with the Proteomics & Peptide Synthesis Core, University of Michigan. Lysate of four independent batches of OT-II Ectosomes and parental OT-II T cells were size-fractionated by SDS-PAGE using a 10% Bis-Tris NuPage Mini-gel (Invitrogen) with the MES buffer system. The mobility region was excised into 20 equally sized bands, which were then digested with trypsin in-gel using a ProGest robot (Digilab). The trypsin digestion process involved the following steps: 1) washing with 25mM ammonium bicarbonate followed by acetonitrile, 2) reduction with 10mM dithiothreitol at 60°C followed by alkylation with 50mM iodoacetamide at room temperature, 3) digestion with sequencing grade trypsin (Promega) at 37°C for 4 hours, and 4) quenching with formic acid. The supernatant containing the digested peptides was analyzed directly without further processing.

Each digest sample was analyzed by nano-MS/LS using a Waters NanoAcquity HPLC system interfaced to a Thermo Fisher Fusion Lumos mass spectrometer. Half of each digested sample was loaded on a trapping column and eluted over a 75µm analytical column at a flow rate of 350nL/min. Both columns were packed with Luna C18 resin (Phenomenex). The mass spectrometer was operated in data-dependent mode, with the Orbitrap operating at 60,000 FWHM (Full Width at Half Maximum) and 15,000 FWHM for MS and MS/MS, respectively. The instrument was run with a 3-second cycle for MS and MS/MS. A total of 10 hours of instrument time was used for the analysis of each sample.

The data obtained from the mass spectrometry analysis were searched using a local copy of Mascot (Matrix Science) with the following parameters: Enzyme: Trypsin/P; Database: SwissProt Mouse (concatenated forward and reverse plus common contaminants); Fixed modification: Carbamidomethyl (C); Variable modifications: Oxidation (M), Acetyl (N-term), Pyro-Glu (N-term Q), Deamidation (N/Q); Mass values: Monoisotopic; Peptide Mass Tolerance: 10 ppm; Fragment Mass Tolerance: 0.02 Da; Max Missed Cleavages: 2. The Mascot DAT files were then parsed into Scaffold software (Proteome Software) for validation, filtering, and to create a non-redundant list per sample. The data were filtered using a 1% protein and peptide false discovery rate (FDR), and a minimum of two unique peptides per protein were required for identification. *T*-test were performed after FDR correction to calculate P values using Normalized Spectral Abundance Factor (NSAF) (*81*) (NSAF = (SpC/MW) / Σ(SpC/MW)N, where SpC represents the spectral counts, MW is the protein molecular weight in kDa, and N is the total number of proteins). $P<0.05$ and fold change >1.8 was used as cutoff to identify enriched proteins in ectosomes.

**Nitrophenyl (NP)-protein conjugates**

To generate hapten NP (4-hydroxy-3-nitrophenyl acetyl) -protein conjugates, OVA (ovalbumin, Sigma), HEL (hen egg lysozyme, Sigma), BSA (bovine serum albumin, fraction V, Sigma), and GFP-GP$^{61\text{-}80}$ were mixed overnight with NP-OSU (4-Hydroxy-3-nitrophenylacetic acid succinimide Ester, Santa Cruz). Insoluble aggregates and particles were removed by centrifugation (10,000*g*×30min), and free NP-OSU was washed out using an Amicon spin-filter (Sigma). Subsequently, the NP-protein conjugates were resuspended in PBS. Based on $OD^{430nm}$ (NP absorption) and $OD^{280nm}$ (protein absorption) detected by a Nanodrop 2000 spectrophotometer (Thermo scientific), NP-protein conjugates were found to have NP/protein ratios at 2.5-4.5. Concentrations of each NP-protein conjugate for antigen-loading B cells were determined by titrating with their ability to upregulate CD69 on QM-B cells. Concentrations of the different NP-protein conjugates that resulted in comparable CD69 upregulation were used for downstream experiments.

**Flow-cytometry**

For analysis by flow-cytometry, live cells were suspended in ice-cold PBS/2% FBS buffer at $0.5\times10^6$ cells/ml, labelled with Fixable Viability Dye eFluor™ 506 (Invitrogen), fixed with 2% PFA for 10 minutes, Fc-blocked with anti-CD16/CD32 (Biolegend) for 30 minutes on ice, and stained with fluorescently labeled antibodies for 30 minutes on ice. For the detection of intracellular transcriptional factors, cells were fixed and permeabilized using Foxp3/Transcription Factor Staining Buffer Set (Invitrogen). The stained cells were washed and filtered through 70µm cell strainer before being analysed on a BD LSRFortessa, equipped with 405nm laser line and associated emission filter sets (450/25nm, 595LP+610/10nm, 655LP+670/15nm, 505LP+525/50nm), a 640nm line with associated emission filters sets (670/15nm, 750LP+780/30nm, 685LP+730/23nm), a 561nm line with associated emission filter sets (586/8nm, 595LP+610/10nm), and 488nm line with associated emission filter sets (690LP+710/25nm, 550LP+586/8nm, 750LP+780/30nm, 635LP+670/7nm, 505LP+530/15nm) Using Flowjo10 software, fluorescently labelled live cells were analysed by serial gates, including size gate of lymphocyte using FSC-A/SSC-A channel, singlets gate using FSC-H/FSC-W and SSC-H/SSC-W channels and live cell gate. Each sample has three replicates. Antibodies used are listed in Table S1.

To detect TCR transfer in T-B cell co-cultures by flow-cytometry, transgenic NP-specific QM-B cells were pulsed with 200µg/ml NP-protein conjugates in the presence of 100ng/ml mIL-4 (Biolegend) for 16 hours, washed, and co-cultured with OT-II Th2 cells at 1:1 ratio or 50:1 ratio for 1-24 hours. After viability dye staining and fixation with 2% PFA, cells were Fc-blocked for 30 minutes on ice, and labeled with fluorescently conjugated antibodies targeting CD3ε, $I\text{-}A^b$, B220 or CD19 on ice for 30 minutes to distinguish T cells and B cells. To detect TCR transfer to B cells, the live singlet cells were gated, TCR transfer to B cells was quantified as percentage of singlet events in the $I\text{-}Ab^{hi}$/ $CD3\varepsilon^+$ region in T-B cultures, defined using unpulsed B cells, to demarcate T and B cell populations. A similar protocol was used to detect TCR transfer in cocultures of ESCRT KO and control T cell, or Tfh cells, but T cells and B cells were cocultured at 1:1 ratio and assayed at 24 hours.

**Cell culture & sorting for SIM imaging**

To prepare cells for SIM imaging, transgenic NP-specific QM-B cells were pulsed with 200µg/ml NP-protein conjugates with 100ng/ml mIL-4 for 16 hours, washed, and co-cultured with OT-II Th2 cells at 1:1 ratio for 24 hours. Cells were then fixed with 4% PFA for 15 minutes, permeabilized with 0.1% Saponin for 5 minutes, quenched with 100mM glycine for 1 hour, Fc-blocked with anti-CD16/32 for 1 hour, and blocked with 5% BSA/PBS for 1 hour. Cells were then stained with in-house-conjugated fluorescently-labeled antibodies in 2% BSA/PBS overnight at

4°C with the mixture of: Alexa-488-anti-I-$A^b$ antibody (AF6-120.1, F/P ratio=3.8) and Alexa-647-anti-CD3ε antibody (17A2, F/P=5.6) for flow-cytometry analysis.

To visualize flow-cytometry-sorted T and B cells using Structured Illumination Microscopy (SIM), antibody-stained cells were post-fixed with 2% PFA and 0.1% glutaraldehyde, and sorted on a Sony SH800 instrument, employing the gating strategy shown in fig. S4B. Briefly, cells were first size-gated using FSC-A/SSC-A channels and subsequent stringent singlet gate applied based on FSC-H/FSC-W and SSC-H/SSC-W channels. The separately cultured OT-II T cells and QM-B cells were used to demarcate 'I-$Ab^-$ TC' and '$TCR^-$ BC' respectively in I-Ab/CD3ε scatter plot. In the T-B coculture, the cell population that has comparable levels of I-Ab with '$TCR^-$ BC' but higher levels of TCR were gated as '$TCR^+$ BC', the cell population that has similar levels TCR to 'I-$Ab^-$ TC' but higher levels of I-Ab, was gated as 'I-$Ab^+$ TC'. Cells were sorted into cold PBS at low flow rates, pelleted, deposited onto poly-L-lysine-treated coverslips, and mounted with ProLong™ Glass Antifade Mountant containing blue-fluorescent DNA stain NucBlue (Hoechst 33342) (Invitrogen).

**Structured illumination microscopy (SIM)**

SIM imaging was performed using a Nikon N-SIM system. Imaging was performed using a 100 × 1.49NA objective (SR HP Apo TIRF 100×H) and an ORCA Flash 4.0 (Hamamatsu) sCMOS camera. Fluorescent antibody-stained cells mounted in sealed coverslips were imaged using a 405nm laser line with 452/22.5nm emission filter, a 488nm line with 525/25nm emission filter, a 561nm line with 593/23nm emission filter, and a 647nm line with 700/37.5nm emission filter, using a software-controlled 3D-SIM acquisition regime. Raw images of each fluorescence channel were recorded as non-saturated 16-bit images with a resolution of 5120×3072 pixels (0.07μm/pixel). The z-step for image acquisition was set to 200nm for 3D image acquisition. Acquired images were reconstructed using the "N-SIM Stack Reconstruction" tool integrated into NIS-Elements software (Nikon). This reconstruction process yielded 16-bit 2048×2048 pixels (0.03μm/pixel) images.

**Crispr-Cas9-mediated gene deletion**

The gRNA expressing vector pKLV2-U6gRNA5(BbsI)-PGKpuro2ABFP, which co-expresses gRNA, a puromycin resistance gene, and blue fluorescence protein (BFP), was a gift from Dr. Kosuke Yusa (Addgene plasmid # 67991(*82*). The gRNA sequences were designed using CHOPCHOP v3 (*83*). gRNA sequences are shown in Table S2. Synthesized oligos (IDT) were annealed and cloned into BbsI-linearized pKLV2-U6gRNA5(BbsI)-PGKpuro2ABFP using T4 ligase (NEB). The resulting construct, along with the second-generation lentiviral package plasmids psPAX2 (Addgene 12260) and pMD2.G (Addgene 12259), was transfected into HEK293T cells using Lipofectamine 3000 (Invitrogen). Virus-containing medium (VCM) was collected 48 hours later for downstream T cell transduction.

To generate recipient T cells, OT-II transgenic mice were crossed with Cas9 knock-in mice to generate F1 progeny that express Cas9 in OT-II T cells. MACS-purified Cas9+ OT-II T cells were stimulated with plate-coated anti-CD3ε/anti-CD28 (5μg/ml each) in Th2 polarization medium (also see ***T cell differentiation and ectosome production***) for 24 hours, prior to incubation with VCM containing 5μg/ml polybrene. The transduced cells were selected using 5μg/ml puromycin for 5 to 7 days. Dead cells were removed using EasySep Dead Cell Removal Kit (Stemcell). The transduction efficiency was confirmed by assessing BFP expression using flow-cytometry, and the gene knockout efficiency was determined by western blot.

**Flow-cytometric intracellular $Ca^{2+}$ measurement**

QM-B cells were pulsed with 100μg/ml of NP-OVA or NP-HEL with 100ng/ml mIL-4 for 48 hours. Cells were washed and loaded with 1μM Indo-1 in serum-free medium and incubated at 37°C for 30 minutes. After several wash steps to remove free dye, the cells were rested in complete medium for an additional 30 minutes. Indo-1-loaded cells were transferred into HBS/HSA imaging buffer (20 mM Hepes, 137 mM NaCl, 5 mM KCL, 0.7 mM Na2HPO4, 6 mM D-Glucose, 1 mM CaCl2, 2 mM MgCl2 and 1% HSA, pH 7.4) at a density of $2\times10^6$ cells/ml. Flow-cytometry was performed using a Bigfoot Spectral Cytometer (Thermofisher) equipped with a 349nm excitation laser. $Ca^{2+}$-free Indo-1 fluorescence was detected using a 473/15nm filter, and $Ca^{2+}$-bound Indo-1 fluorescence was detected using a 420/10nm filter. A heated tube holder maintained samples at a temperature of 37°C throughout experiments. For each sample, an initial 30-second baseline measurement was collected. After adding the indicated stimuli, B cell Indo-1 fluorescence was monitored for 3-4 minutes. The ratio of $Ca^{2+}$ bound/ $Ca^{2+}$ free fluorescence was analyzed using FlowJo 10 and plotted in GraphPad Prism 9.

**Enzyme-linked immunosorbent assay (ELISA)**

To detect mIL-2 and mIL-4 in the culture supernatant, sandwich ELISA was performed. The assay was performed using 1µg/ml capture antibody and 1µg/ml corresponding biotinylated detecting antibody, followed by labeling with ExtrAvidin-Peroxidase (Sigma E2886, 1:1000 dilution) and the wells developed using 1-Step™ Ultra TMB-ELISA Substrate Solution (Thermo Scientific™). Sandwich ELISA was also used for detection of mouse IgG. IgG levels were quantified using pan-IgG capture antibodies, used at a coating concentration of 1µg/ml, and antigen-specific IgG was captured using the respective antigens (NP-BSA/NP-HEL/OVA/HEL) adsorbed in wells at a coating concentration of 100µg/ml. Cytokine and antibody concentrations were determined from calibration curves of doubling-dilutions of known concentration of reference preparations of cytokines or antibodies. All antibody pairs and calibration references used in the assays were listed in ***Table S1***.

**Mouse vaccination and memory-enriched B cell (MEBC) phenotyping**

To induce memory B cells in 8–10-week-old $CD45.2^+$ C57bl/6 mice, mice were intraperitoneally injected with an emulsion containing 100μg NP-BSA antigens in 50μl PBS and 50μl Complete Freund's Adjuvant (CFA, Sigma). For *in vitro* experiments, spleen B cells were isolated from hyperimmunized mice following 4 secondary inoculations using incomplete Freund's Adjuvant (IFA, Sigma) instead of CFA, and maintaining a 2-week interval between vaccinations.

Vaccination response and MEBC induction was monitored by measuring anti-NP antibodies in the serum, and by identifying $NP^+$ B cells with a memory phenotype, using 'spiked in' lymphocytes from age-matched unvaccinated mice as negative controls, as described in detail elsewhere (*36*). Briefly, splenocytes of $CD45.2^+$ C57bl/6 mice vaccinated with NP-BSA were mixed with splenocytes from age-matched unvaccinated $CD45.1^+$ C57bl/6 mice. The resulting mixed lymphocytes were processed for antibody staining in a single staining tube to allow for the simultaneous identification of NP-specific BC (Live $CD19^+CD38^+PNA^-CD45.2^+NP^+$) from vaccinated mice, and their naïve counterparts (Live $CD19^+CD38^+PNA^-CD45.1^+CD23^+CD21/35^+$) from unvaccinated $CD45.1^+$ mice. The gating strategy is as shown in fig. S5A.

To monitor the Ig class-switch of NP specific memory-enriched B cells (MEBC), CD81, a MEBC maker identified in (*36*) and confirmed in our experiment fig. S5B, was used to identify $NP^+$ MEBC (Live $CD19^+CD38^+CD81^+NP^+$), the gating strategy is as shown in fig. S5A. Surface expression of IgM, IgG and IgD on $NP^+$ MEBC was determined by Flow cytometry.

**Tfh cell induction and TCR transfer *in vivo***

8–10-week-old WT C57bl/6 mice were *i.p.* vaccinated with an emulsion containing 100μg OVA in 50μl PBS and 50μl CFA. To increase pOVA/I-Ab specific T cell precursor, freshly isolated OTII T cells were adoptively transferred *i.v.* to WT C57bl/6 by tail vein injection one day before immunization. One week post vaccination, splenocytes were harvested and processed for flow-cytometry analysis. After gating live singlets (as shown in fig. S7A), B cells ($CD19^+$) with transferred TCRβ were analysed and quantified. Tfh differentiation of pOVA/I-Ab specific CD4 cells was determined by expression of Tfh markers (PD1, CXCR5, ICOS, CD40L and BCL6) and lack of CCR7 expression. The cell counts were quantified by a spike-in CountBright Plus Absolute Counting Beads (Invitrogen).

To isolate pOVA/I-Ab specific Tfh cells for *in vitro* coculture experiments, WT C57bl/6 mice seeded with 1 million OTII T cells were immunized as described above, one-week later splenic CD4 T cells were enriched by negative MACS, as shown in ***Cell isolation and purification***. Pooled CD4 T cells were stained with fluorescently conjugated pOVA/I-$A^b$ tetramer and CXCR5/PD-1 antibodies, and triple-positive cells were sorted on a Sony SH800 instrument. After 4 hour rest in complete RPMI 1640, Tfh cells were co-cultured with antigen pulsed QM-B cells to assay TCR transfer *in vitro*, following the steps described in ***Flow-cytometry***.

**Somatic hypermutation assay (SHM).**

8-10-week-old $Rag1^{-/-}$ mice were *i.v.* adoptively transferred with OT-II T cells and QM-B cells ($2\times10^6$ each), and *s.c.* vaccinated with 100μg NP-OVA mixed with CFA adjuvant at tail base after 3 days. Lymphocytes from the inguinal lymph node were collected on day 4 post-vaccination for flowcytometry assay. To evaluate SHM differences, $TCR^+$B cells and $TCR^-$B cells were separated by sorting using the Sony H800 instrument. Genomic DNA was purified from the sorted $TCR^+$B cells, $TCR^-$B cells, and freshly isolated untreated QM B cells (as a negative control) using the Wizard® Genomic DNA Purification Kit (Promega). A previously reported method was used to analyse the BCR mutations (*20, 84*). Briefly, a semi-two-step PCR was performed to amplify and introduce assembly overhangs to the VDJ region of QM-BCR VH17.2.25 using the primers: 1st step: 5'-TTCAGAGGTTCAGCTGCAGCAGT-3' and 5'-CTYACCTGAGGAGACDGTGA-3'; 2nd step: 5'-GATCCCGGTACTCGAGTTCAGAGGTTCAGCTGCAGC-3' and 5'-TCTAGAGTCGCGGCCGCGAGGAGACGGTGACTGAGG-3'. The Phusion High-Fidelity Enzyme with a low error rate of $4.4\times10^{-7}$ (Thermo Scientific™) was used for PCR amplification. The resulting PCR products were inserted into XhoI/NotI linearized pHR-sin vector using HD-infusion cloning (Takara) and then transfected into One Shot™ OmniMAX™ 2 T1R Chemically Competent E. coli (Invitrogen). Plasmids were subsequently miniprepped from individual colonies and subjected to Sanger sequencing. The obtained sequences were aligned and compared using MegAlign Pro (DNASTAR). The mutation rates of nucleotides were plotted using GraphPad Prism 9.

***In vitro* help assay**

Splenic B cells from QM transgenic mice or NP-BSA-hyper-immunized wild-type C57bl/6 mice were pulsed with NP-protein conjugates and 100ng/ml mIL-4 overnight in complete medium. After washing, B cells were cultured with ectosomes derived from the indicated numbers of T cells (cultured for 48 hours), sEcto ($2.7\times10^{10}$), or OT-II T cells with CRISPR-edited TSG101/VPS4 knockouts, along with 100ng/ml mIL-4 for 1 week. NP-specific or total IgG in the supernatant was detected using ELISA.

**Adoptive transfer model for T-B collaboration**

To test ectosome-mediated help *in vivo*, an adoptive transfer model of T cell-mediated B cell help was established using Rag1$^{-/-}$ mice. Splenic B cells, isolated 8 weeks after C57bl/6 mice were vaccinated with 50μg OVA in CFA, were loaded overnight with 0.1mg/ml OVA *in vitro*, and $10^6$ thoroughly washed B cells were transferred intravenously to Rag$^{-/-}$ recipients, along with $2\times10^{10}$ OTII ectosomes, or PBS vehicle as a negative control. Serum samples were collected three weeks after the transfer of B cells and ectosomes, and analyzed for OVA-specific IgG antibodies by ELISA.

**Aged mouse vaccination**

To evaluate the effect of ectosomes on vaccine-induced antibody responses in aged mice, aged (75-78-week old) C57bl/6 mice were subcutaneously injected with 50μl of PBS containing 50μg OVA, 50μg HEL, and 1μg CPG ODN 2395 (InvivoGen). One hour later, the mice were randomly divided into groups and intravenously injected with either $8\times10^{10}$ OT-II ectosomes or $8\times10^{10}$ synthetic T cell mimetic ectosomes (sEcto) (see also ***Synthetic ectosomes***). In some experiments, adult mice (18 week-old) were included for comparison. Serum samples were collected at the specified time points, and antibodies were measured using ELISA.

**Synthetic ectosomes**

Liposomes containing 80 mol% 18:1 (Δ9-Cis) PC (DOPC), 10 mol% 18:1 PE-NBD, and 10 mol% 18:1 PE-benzylguanine (Avanti Polar Lipids), were prepared by extrusion through a 100nm pore-size as previously described (*85, 86*). Lipids in chloroform were transferred to 15ml glass test-tubes prewashed with chloroform and filled with argon to prevent oxidation. The chloroform was evaporated using a gentle N2 gas stream while warming in a 37°C water bath for 10 minutes. The resulting lipid residue was then lyophilized under argon for 2 hours, and resuspended in degassed liposome buffer (25mM Tris, 150mM NaCl, pH 8.0, 0.2μm filtered, nitrogen gas-treated) at a final total lipid concentration at 4mM. The resuspended lipids were extruded through a 100nm pore-size polycarbonate membrane (Whatman) using a glass Mini-Extruder (Avanti Polar Lipids) and two gas-tight syringes (1000μl, Avanti Polar Lipids). After 12 rounds of extrusion, the formed liposomes were diluted to 0.4mM in liposome buffer, filtered through a sterile 0.2μm filter, aliquoted to under argon, and stored at 4°C. PE-NBD lipids were shielded from light at al stages. PE-NBD was omitted from the lipid mix when fluorescent liposomes were not required. No functional differences were found between liposomes with or without PE-NBD. The diameter and density of the liposomes was measured by NTA as described in ***Nanoparticle tracking analysis (NTA)***, after 5,000x dilution of the 0.4mM stock in filter-degassed PBS.

To conjugate liposome with SNAP-tagged proteins, an empirically determined concentrations yielding the desired surface levels of SNAP-containing proteins on liposomes were used. Briefly, 150μl 0.4mM liposomes was mixed with 30μg/ml SNAP-tagged proteins in 1ml PBS at 37°C for 5 hours. The unbound proteins were washed out through UV-irradiated 100KD Amicon centrifugal filters (Sigma) for three times. The protein conjugated liposomes were resuspended in 150μl of PBS or culture medium. To check and quantify the protein conjugated on liposomes, SNAP-tagged proteins were first conjugated with Alexa-647 (Alexa-647-SNAP, F/P=1.55; Alexa-647-OT-II scTCR-SNAP, F/P=3.5; Alexa-647-CD40L-SNAP, F/P=6.2). The liposomes with Alexa-647-conjugated proteins were made as described before and resuspended in filtered-degassed PBS for detection using Ze5 flow-cytometer as described in ***Small-particle flow-cytometry***, but instead of being stained with PKH67 as for the ectosomes, liposomes were discriminated by the fluorescent signal from the incorporated PE-NBD (Ex/Em=460/535nm). Protein mass in sEcto were quantified using Pierce™ Detergent Compatible Bradford Assay Kit

(Thermo Scientific), following manufacturer's instruction, and the mass equivalent liposome-free proteins were used to compare with sEcto to help B cells *in vitro* (fig. S8D) and *in vivo* (fig. S8E).

**Bead bilayer**

To validate the binding of SNAP-proteins with specific antibodies or tetramers, and to quantify the molecular density of proteins attached on bilayers, a flow-cytometry experiment was conducted on bead bilayers. Silica microspheres (1µL, SS06N, Bangs Laboratories, Inc) were mixed with liposomes (4µL) having the same composition as SLB and incubated for 10min at room temperature. The blocking and SNAP-protein loading processes were performed similarly to SLB experiments but in eppendorf tubes. Following the loading process, samples were washed by centrifugation in HBS/HSA buffer. Antibody or tetramer staining and data acquisition were carried out using the same cell-based flowcytometry protocol described earlier.

**Supported lipid bilayers**

Supported lipid bilayers (SLBs) were formed by depositing liposomes onto piranha-cleaned glass coverslips in Bioptechs flow chambers, as previously described (*85*). The liposomes composition was 12.5 mol% DGS-NTA and 7.5 mol% 18:1 PE-benzylguanine. Bilayers formed on coverslips were blocked with a 4% casein solution containing 10mM $NiSO_4$, mICAM-1-His10 was attached to the bilayers using $Ni3^+$-charged NTA-DGS, while scOT-II TCR-SNAP fusion proteins or SNAP-tag alone was attached by conjugation with benzylguanine headgroups. The bilayers had an approximate density of 200 molecules/$\mu m^2$ of SNAP-tag or TCR-SNAP and 350 molecules/$\mu m^2$ of ICAM-1.

**Intracellular $Ca^{2+}$ imaging**

Intracellular $Ca^{2+}$ imaging of fluo-4-loaded QM transgenic B cells was performed by epifluorescence microscopy. QM-B cells were pulsed with NP-OVA or NP-HEL overnight and loaded with Fluo-4. Fluo-4-loaded B cells were resuspended in HBS/HSA imaging buffer and introduced into heated Bioptechs flow chambers maintained at 37°C. Glass-supported lipid bilayers (SLBs) containing TCR-SNAP or SNAP and ICAM-1. The B cells were imaged starting from the moment they settled on the scTCR/ICAM-1-containing SLBs using a 20× air objective. Imaging was conducted for 2 minutes using a 488nm laser for Fluo-4 excitation. Fluorescence intensity was quantified using Fiji/ImageJ from acquired 16 bit images, and presented as change in fluorescence ($\Delta F$) relative to the initial fluorescence intensity ($F_0$). The $\Delta F/F_0$ values were plotted over time as a measure of intracellular $Ca^{2+}$ influx.

**Total internal reflection fluorescence microscopy (TIRFM) of B cell synapses**

The synaptic contact interfaces between QM-B cells and SLBs were visualized by TIRFM. $10^6$ NP-OVA or NP-HEL pulsed B cells suspended in HBS/HSA were introduced into heated flow chambers and allowed to interact with TCR/ICAM-1-bearing SLBs formed on coverslip. After 10-minute incubation in flow chambers, B cells interacting with SLBs were fixed using 3% PFA/HBS for 15 minutes. Flow chambers were maintained at 37ºC throughout using a filament heater mount. Following fixation, the cells were permeabilized with 0.1% saponin for 5 minutes, quenched with 100mM glycine for 1 hour, Fc-blocked with anti-CD16/32 for 1 hour, and blocked with 5% BSA/PBS for 1 hour. Cells were then sequentially stained with antibodies in 2% BSA/PBS in the following order: 1. Alexa-488-anti-I-Ab antibody (AF6-120.1, F/P ratio=3.8, in-house conjugated), 1 hour, 2. Rabbit-anti-phospho-Tyrosine (P-Tyr-1000, Cell Signaling Technology), 1 hour, 3. Alexa-647-F(ab')2-Goat anti-Rabbit (Invitrogen), 1 hour. All incubation and wash steps were performed within flow chambers. B cell synaptic contacts were imaged by SRIC and TIRFM, with a TIRF excitation field extending ~100nm from the coverslip, on a Nikon-STORM inverted microscope fitted with a NIKON CFI SR HP Apo TIRF 100×objective (NA 1.49) and EMCCD camera (DU-897U, Andor). Alexa-488 was imaged by 488nm laser with 525/25nm filter, and Alexa-647 was imaged by 633nm laser with 700/40nm filter.

**Statistical analysis**

Unpaired T-test, paired T-test and One-way ANOVA (with correction for multiple comparisons) were performed to compare multiple sample means using GraphPad PRISM9 software.

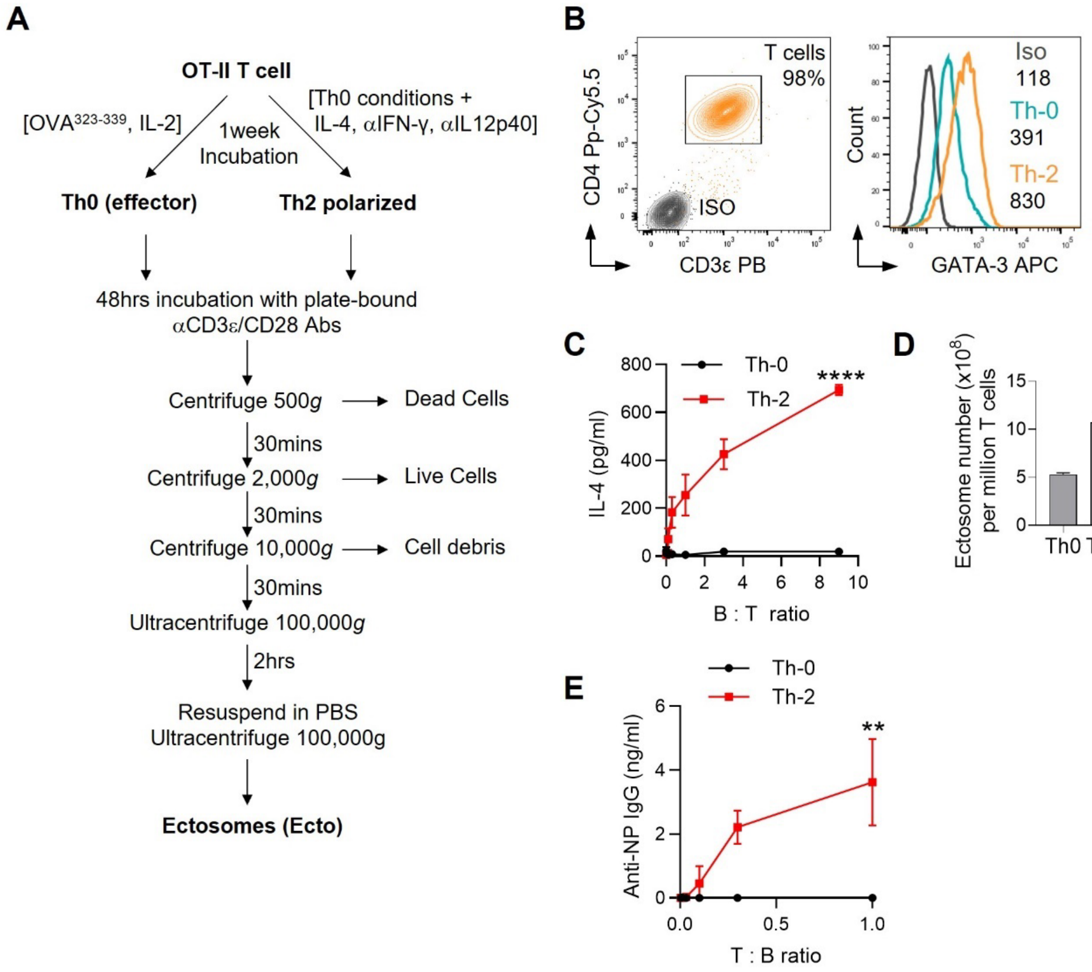


**Fig. S1. Ectosome isolation and helper activity of Th2-polarized and Th0 OTII T cells.** (**A**) Schematic of ectosome production and isolation from activated Th2-polarized OT-II T cells. Ecto, ectosomes. (**B**) Flow-cytometry of GATA-3 transcription factor levels in Th2 polarized and Th0 effector OT-II T cells, numbers under legends are the corresponding mean fluorescence intensities (MFI) in arbitrary unit. (**C**) Measurement of IL-4 release by Th0 and Th2-polarized OT-II T cells ($10^4$/well) in cocultures with QM B cells. QM B cells were pre-incubated overnight with 100µg/ml NP-OVA and 100ng/ml IL-4, and washed 4 times before coculture with OT-II T cells at the indicated ratios for 24 hrs, and the IL-4 measured in culture supernatants by ELISA. Data are representative of 3 independent experiments. (**D**) Nanoparticle tracking analysis (NTA) quantitation of vesicle release by Th0 and Th2 OT-II T cells activated with immobilized anti-CD3ε/CD28 antibodies. Mean ± s.e.m. is indicated. (**E**) Th0 or Th2-polarized OT-II T cells and NP-OVA-pulsed B cells ($5 \times 10^4$/well) were incubated for 7 days at the indicated ratios under culture conditions as in (C). Culture supernatants were then harvested and assayed for released anti-NP IgG antibodies by ELISA. Data are representative of 3 independent experiments. Means were compared by two-tailed *t*-test. In (C) data points corresponding to B:T ratio=9 were compared, and in (E) data points corresponding to T:B ratio=1 were compared. ****, $P < 0.0001$; **, $P < 0.01$.

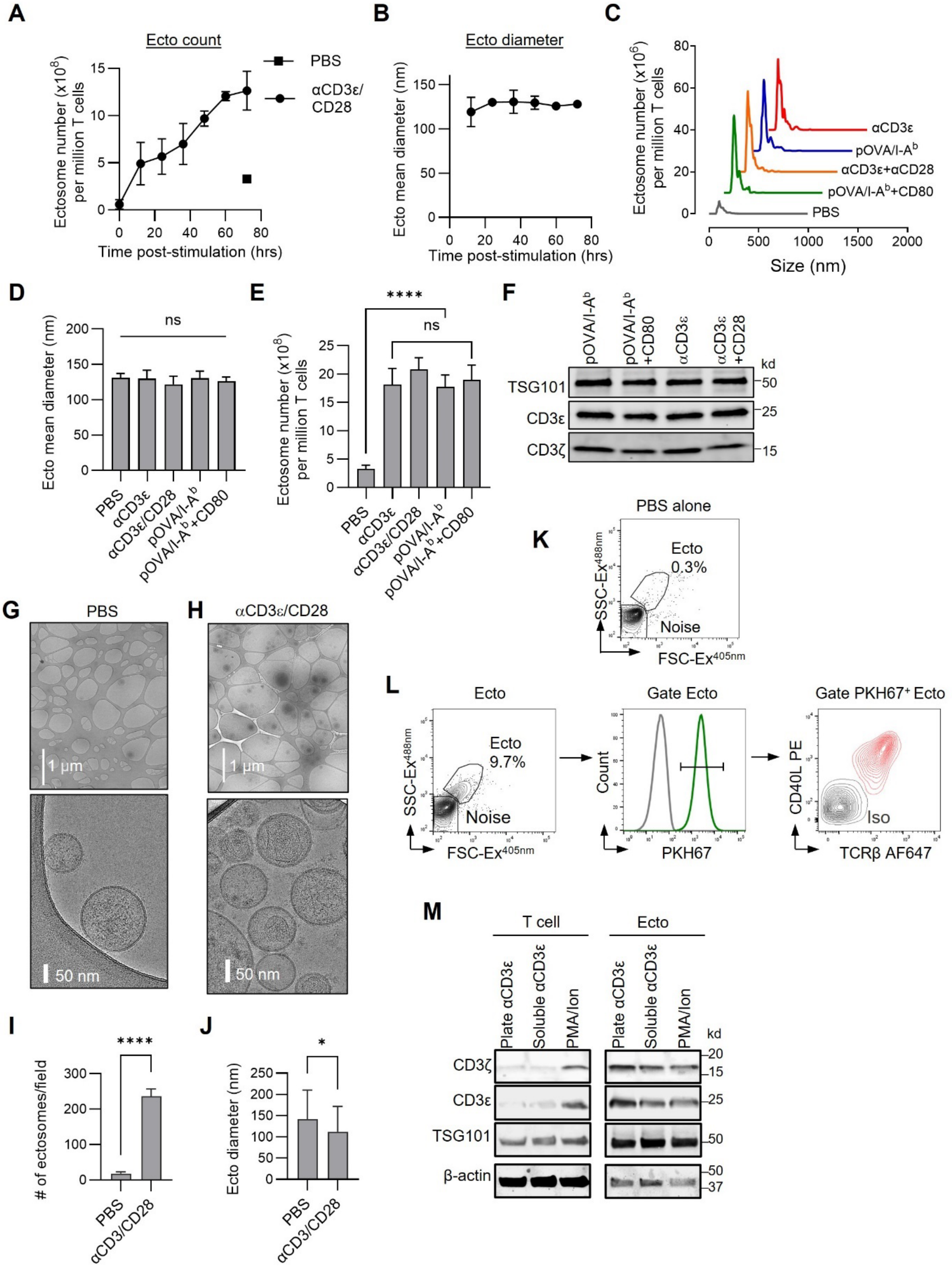


**Fig. S2. Biophysical and biochemical characterization of ectosomes released by OT-II T cells.** (**A**) Timeseries of ectosome release in culture supernatants of OT-II T cells activated in anti-CD3ε/CD28 antibody-coated wells. Ectosome numbers were quantified by NTA analysis, and normalized to the number of T cells in wells. PBS, T cells incubated in uncoated wells.(**B**)Time series of mean ectosome diameter measured by NTA. (**C**) Count and size profile of ectosomes from T cells left unstimulated (PBS), or activated with anti-CD3ε antibodies, with or without anti-CD28 antibodies, or recombinant pOVA/I-$A^b$ monomers, with or without recombinant CD80. All proteins were adsorbed onto plastic culture wells. Ectosomes were collected and isolated from culture supernatants after 72hrs incubation. Mean ± s.e.m is shown. Data are representative of three independent experiments. (**D**) Quantitation of mean diameter of ectosomes released by T cells activated by the indicated stimuli as in (C). (**E**) Quantitation of number of ectosomes released by T cells activated by the indicated stimuli. (**F**) Immunoblot of TCRζ and TSG101 in ectosomes (Ecto) or parental T cells activated with anti-CD3ε/CD28. Lanes were loaded with 3.2 μg total protein/lane. Data are representative of five independent experiments. (**G, H**) Transmission electron cryomicroscopy (cryo-EM) projection images of ectosomes in vitreous ice. Ectosomes were isolated from supernatants of OT-II T cells activated for 48hrs with anti-CD3ε/CD28 antibodies adsorbed to plastic culture wells in PBS (G), or wells were treated with PBS alone (H); (Top panels) low magnification images of ectosomes in vitreous ice, located in the voids of lacey carbon films; (Bottom panels) are high magnification images of (Top panels). (**I**) Quantitation of ectosomes released from unstimulated and antibody-activated T cells, normalized as number of vesicles/field. **(J)** Quantitation of vesicle diameters. Graph bars show means and standard deviations from four randomly chosen fields or from 50 randomly chosen vesicles (**K**) Small particle flow-cytometry of a blank PBS-only sample showing background noise and the ectosome size gate (Ecto). (**L**) Distribution of TCR and CD40L levels on ectosomes measured by small-particle flow-cytometry. (Left panel) scatter and size characteristics of ectosomes; (middle panel) ectosomes labeled with green-fluorescent lipid dye PKH67 (green) and unlabeled ectosomes (gray); (right panel) contour plot of TCRβ and CD40L levels on $PKH67^+$ ectosomes (red) and ectosomes labeled with isotype control antibodies for both markers (black). (**M**) Immunoblot of TCR and Tsg101 in ectosomes, and parental OTII Th2 cells. Each lane was loaded with 10μg total protein. T cells were stimulated *via* the TCR using plate-bound anti-CD3ε antibody (Plate αCD3ε) or anti-CD3ε antibody in solution (Soluble αCD3ε), or activated without engaging the TCR using PMA/Ionomycin (PMA/Ion). Data are mean ± s.d.. Means were compared using two-tailed *t*-test or one-way ANOVA, corrected for multiple comparisons. ****, $P < 0.0001$; *, $P < 0.05$; ns, not significant ($P > 0.05$).

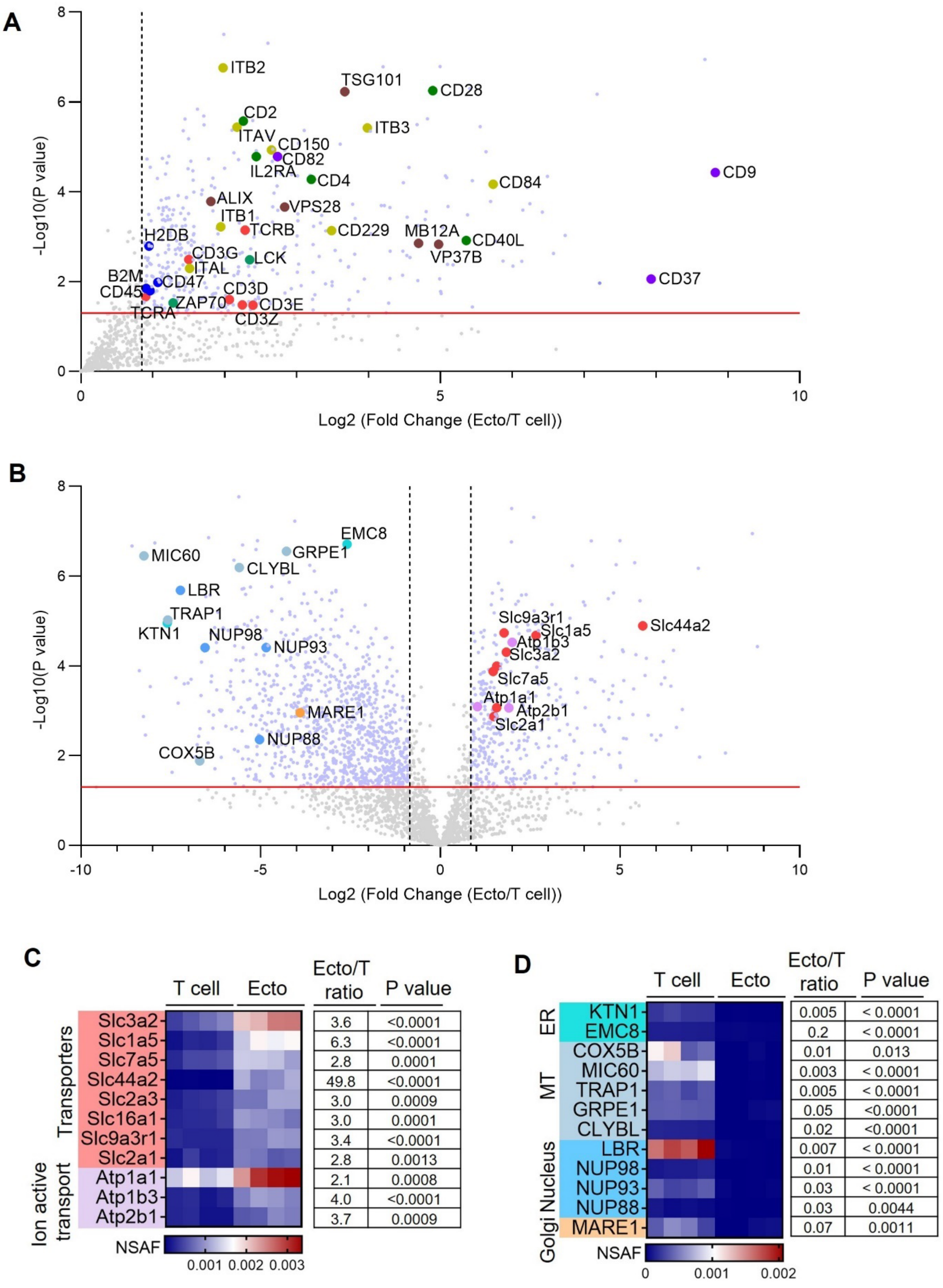
A
-Log10(P value)
Log2 (Fold Change (Ecto/T cell))
ITB2
TSG101
CD28
CD2
ITAV
ITB3
CD150
CD82
IL2RA
CD4
CD84
CD9
ALIX
VPS28
ITB1
TCRB
CD229
MB12A
CD40L
VP37B
H2DB
CD3G
LCK
ITAL
B2M
CD47
CD45
CD3D
CD37
ZAP70
CD3E
TCRA
CD3Z
B
EMC8
MIC60
GRPE1
CLYBL
LBR
TRAP1
KTN1
NUP98
NUP93
Slc9a3r1
Slc1a5
Slc44a2
Atp1b3
Slc3a2
Slc7a5
Atp1a1
Atp2b1
Slc2a1
MARE1
NUP88
COX5B
C
T cell
Ecto
Ecto/T ratio
P value
Transporters
Ion active transport
Slc3a2 3.6 <0.0001
Slc1a5 6.3 <0.0001
Slc7a5 2.8 0.0001
Slc44a2 49.8 <0.0001
Slc2a3 3.0 0.0009
Slc16a1 3.0 0.0001
Slc9a3r1 3.4 <0.0001
Slc2a1 2.8 0.0013
Atp1a1 2.1 0.0008
Atp1b3 4.0 <0.0001
Atp2b1 3.7 0.0009
NSAF
0.001 0.002 0.003
D
T cell
Ecto
Ecto/T ratio
P value
ER
MT
Nucleus
Golgi
KTN1 0.005 < 0.0001
EMC8 0.2 < 0.0001
COX5B 0.01 0.013
MIC60 0.003 < 0.0001
TRAP1 0.005 < 0.0001
GRPE1 0.05 <0.0001
CLYBL 0.02 <0.0001
LBR 0.007 < 0.0001
NUP98 0.01 < 0.0001
NUP93 0.03 < 0.0001
NUP88 0.03 0.0044
MARE1 0.07 0.0011
NSAF
0 0.001 0.002

**Fig. S3 Proteomic analysis of ectosomes released from activated and Th2-polarized OT-II T cells.** (**A**) Volcano plot of proteins enriched in ectosomes. Light purple dots represent proteins above the fold-enrichment cut-off (>1.8) and P value cutoff <0.05 (set for FDR 1% for all proteins), light grey dots represent proteins below the cut-offs, and larger labeled dots represent selected proteins of interest shown in Fig. 1E. (**B**) Full volcano plot showing proteins enriched and depleted in ectosomes. Light purple dots represent significantly enriched or depleted proteins (1% FDR, Fold change > 1.8, P < 0.05), and light grey dots represent proteins not significantly enriched in either ectosomes or T cells. Large dots represent selected proteins of interest. (**C**) Plasma membrane-associated proteins enriched in ectosomes relative to parental T cells. (**D**) T cell endomembrane-associated proteins depleted in ectosomes relative to parental T cells.

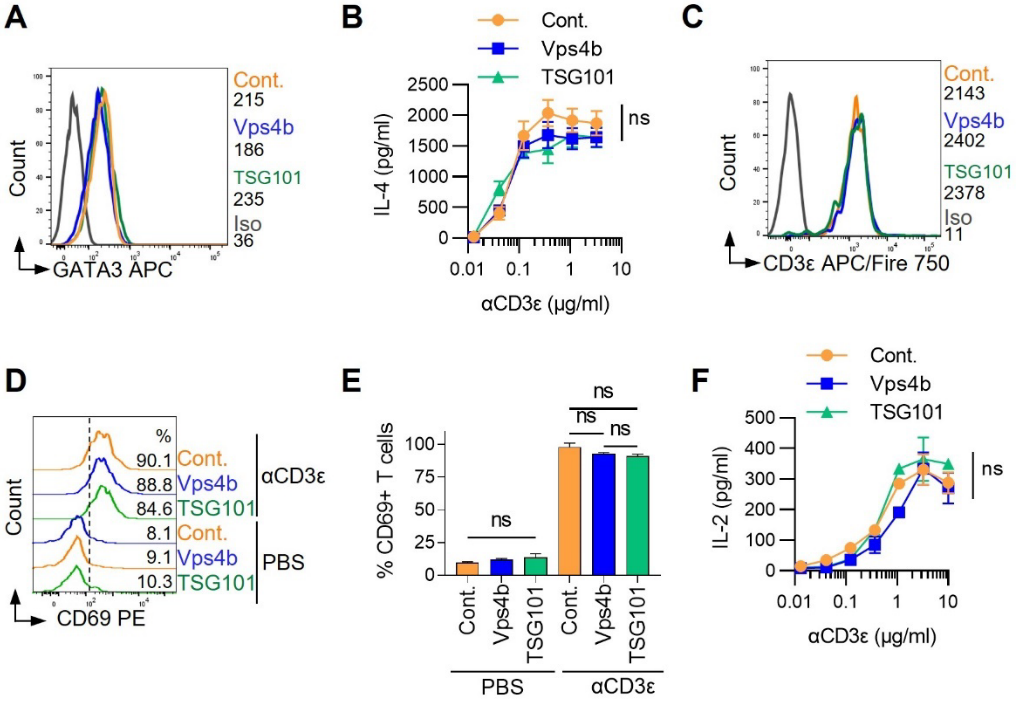


**Fig. S4. Phenotypic and functional characterization of CRISPR-edited TSG101 and VPS4B knockout OTII$^{CAS9}$ T cells.** (**A**) Histogram overlay of GATA3 levels in ESCRT (TSG101 or VPS4b) and control knockout Th2-polarized OT-II$^{CAS9\text{-}GFP}$ T cells. (**B**) IL-4 production by ESCRT (TSG101 or VPS4b) and control knockout Th2-polarized OT-II$^{CAS9}$ T cells. T cells were stimulated with plate-bound anti-CD3ε antibody for 24hrs and culture supernatants harvested for cytokine measurements by ELISA. Data are representative of 3 independent experiments. (**C**) surface TCR levels, in TSG101, VPS4b and control OTII$^{CAS9}$ KO T cells measured by flow-cytometry. ISO, isotype control antibody. Numbers in legend indicate mean fluorescence intensity in arbitrary units. (**D**) CD69 upregulation of the indicated OT-II$^{CAS9}$ T cells following activation with plate-bound anti-CD3ε antibody. Numbers represent %CD69$^{+}$ T cells. Unstimulated T cells (PBS) are shown for comparison. (**E**) Quantitation of CD69 upregulation as in (D). Results are representative of 4 independent experiments. (**F**) IL-2 production by ESCRT (TSG101 or VPS4b) and control knockout Th2-polarized OT-II$^{CAS9\text{-}GFP}$ T cells. T cells were stimulated with plate-bound anti-CD3ε antibody for 24hrs and culture supernatants harvested for cytokine measurements by ELISA. Data are mean ± s.d.. Data are representative of 3 independent experiments. For (B), (E) and (F), means at the 10μg/ml αCD3ε data point were compared by one-way ANOVA corrected for multiple comparisons. ns, $P>0.05$.

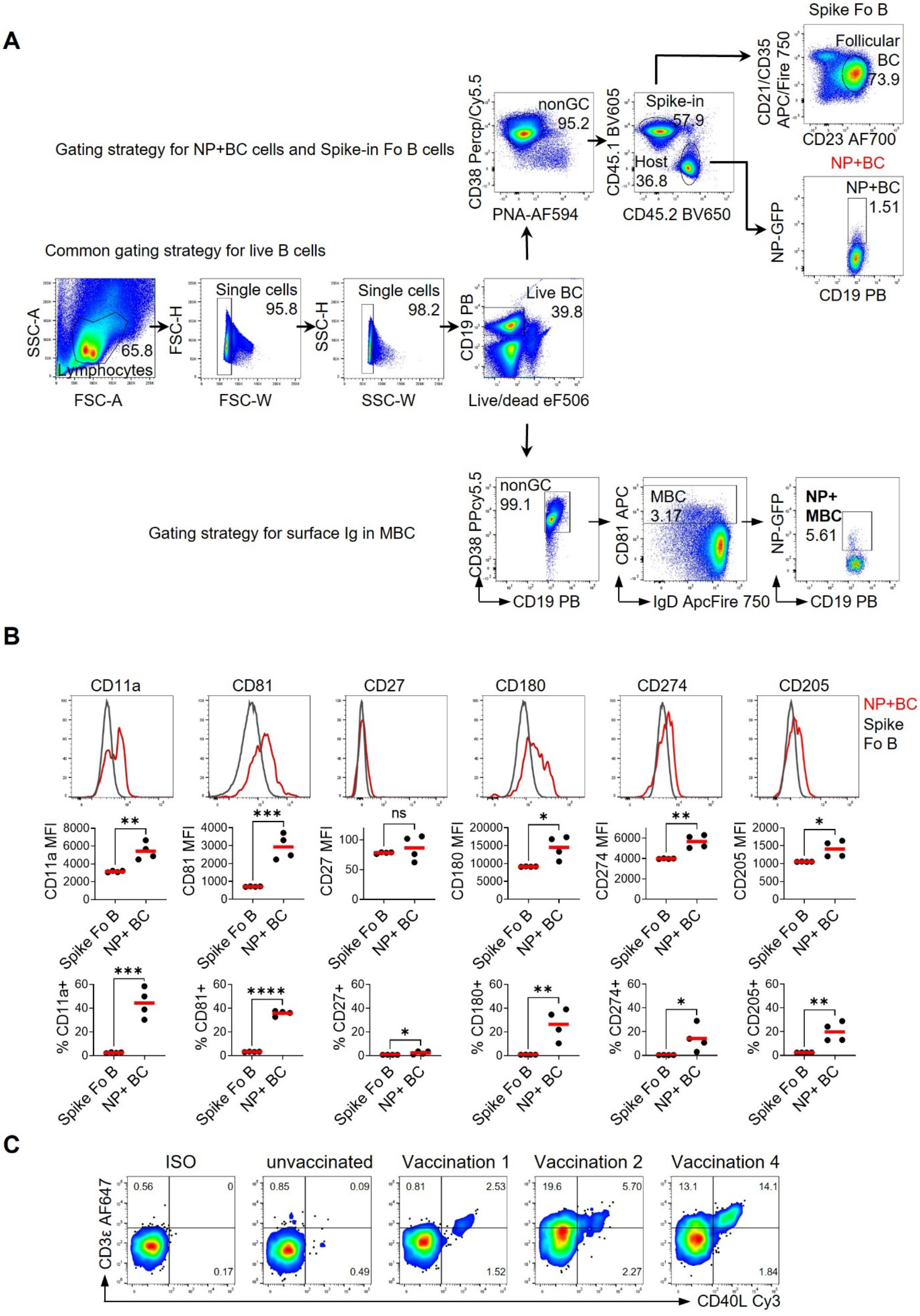

A
Gating strategy for NP+BC cells and Spike-in Fo B cells
Common gating strategy for live B cells
Gating strategy for surface Ig in MBC
Lymphocytes
65.8
Single cells
95.8
Single cells
98.2
Live BC
39.8
nonGC
95.2
Spike-in
57.9
Host
36.8
Spike Fo B
Follicular BC
73.9
NP+BC
NP+BC
1.51
nonGC
99.1
MBC
3.17
NP+ MBC
5.61
SSC-A
FSC-A
FSC-H
FSC-W
SSC-H
SSC-W
CD19 PB
Live/dead eF506
CD38 Percp/Cy5.5
PNA-AF594
CD45.1 BV605
CD45.2 BV650
CD21/CD35 APC/Fire 750
CD23 AF700
NP-GFP
CD19 PB
CD38 PPcy5.5
CD19 PB
CD81 APC
IgD ApcFire 750
NP-GFP
CD19 PB
B
CD11a
CD81
CD27
CD180
CD274
CD205
NP+BC
Spike Fo B
CD11a MFI
CD81 MFI
CD27 MFI
CD180 MFI
CD274 MFI
CD205 MFI
% CD11a+
% CD81+
% CD27+
% CD180+
% CD274+
% CD205+
Spike Fo B
NP+ BC
**
***
ns
*
**
*
***
****
*
**
*
**
C
ISO
unvaccinated
Vaccination 1
Vaccination 2
Vaccination 4
0.56
0
0.17
0.85
0.09
0.49
0.81
2.53
1.52
19.6
5.70
2.27
13.1
14.1
1.84
CD3ε AF647
CD40L Cy3

**Fig. S5. Hyperimmunization of C57bl/6 mice to induce memory-enriched B cells (MBEC).** (**A**) Size and singlet gating, and fluorescence channel gating strategy for phenotypic characterization of splenic B cells isolated from NP-BSA hyperimmunized mice. Splenocytes isolated from age-matched unimmunized CD45.1 C57bl/6 mice were 'spiked' into samples from immunized CD57bl/6 (CD45.2) mice as a source of reference follicular B cells, which are live $CD19^{+}CD38^{+}PNA^{-}CD45.1^{+}CD23^{+}CD21/35^{+}NP^{+}$ B cells from immunized CD45.2 mice were identified as live $CD19^{+}CD38^{+}PNA^{-}CD45.2^{+}NP^{+}$ cells. CD81 was used to identify $NP^{+}$ memory B cells (MBC) defined as live $CD19^{+}CD38^{+}CD81^{+}NP^{+}$. (**B**) Histogram overlays of surface levels (MFI and percentages) of the indicated memory B cell markers in $NP^{+}$ B cells from hyperimmunized mice, and corresponding spiked-in follicular CD45.1 B cells (FoB). Results are pooled from 2 independent experiments. Data points represent individual mice. Means were compared by two-tailed *t*-test. (**C**) Pseudocolored dot-plots, acquired by small particle flow-cytometry, of lipid vesicles in mouse plasma, labeled with fluorescent lipophilic dye PKH67, and with fluorescently conjugated antibodies against TCR (CD3ε) and CD40L. Plasma was collected for analysis 1 week after the indicated serial vaccination with NP-BSA (summarized in the protocol timeline in Fig. 2D). Plasma was pooled from 5 mice for each timepoint. Plots are representative of 3 independent experiments. ***, P < 0.001; **, P < 0.01; *, P < 0.05; ns, not significant (P > 0.05).

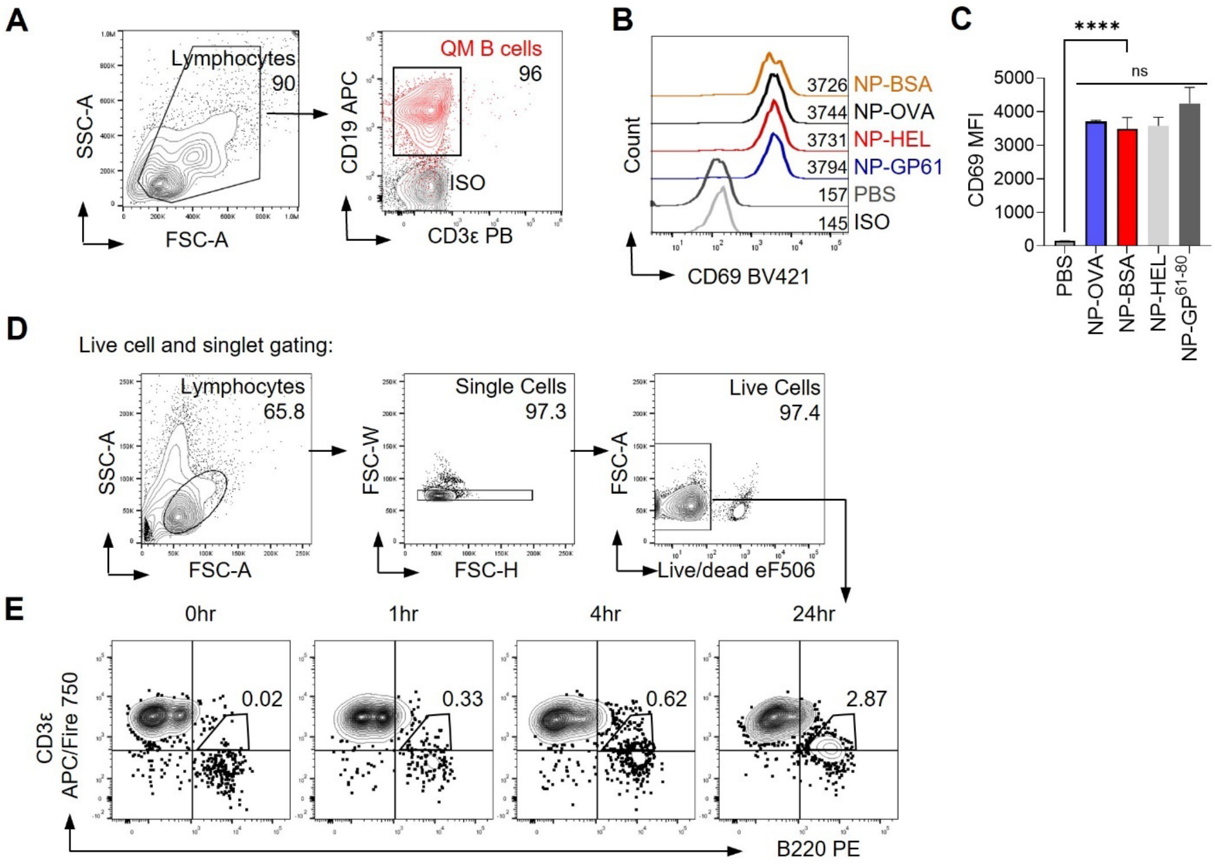


**Fig. S6. Phenotypic and functional characterization of QM transgenic B cell.** (**A**) Representative flow-cytometry contour plots of negatively-enriched NP-specific transgenic QM B cells. (Left panel) Size gate of QM B cells; (Right panel) QM B cells labelled with fluorescent antibodies for B (CD19) and T (CD3ε) cell markers. ISO, antibody isotype control. Numbers within panels indicate percentage of events within indicated regions of interest. (**B**) Representative histogram overlay of NP-specific QM B cell CD69 upregulation, as a measure of activation, analyzed by flow-cytometry. QM B cells were incubated with 200μg/ml of NP-OVA, NP-HEL, NP-BSA, or NP-GP61 for 24hrs. Numbers in legend correspond to mean fluorescence intensity. ISO, isotype control antibody. (**C**) Quantitation of CD69 upregulation as in (B). Results are representative of 4 independent experiments. (**D**), (**E**) Flow-cytometry analysis of cocultured QM B cells, pulsed for 48hrs with 500μg/ml NP-OVA antigens, and Th2 OT-II T cells, at a T:B cell ratio of 50:1 for the indicated times (hrs). Contour plots of live cell singlet gating (**D**) and CD3ε/B220 expression (**E**) are shown.

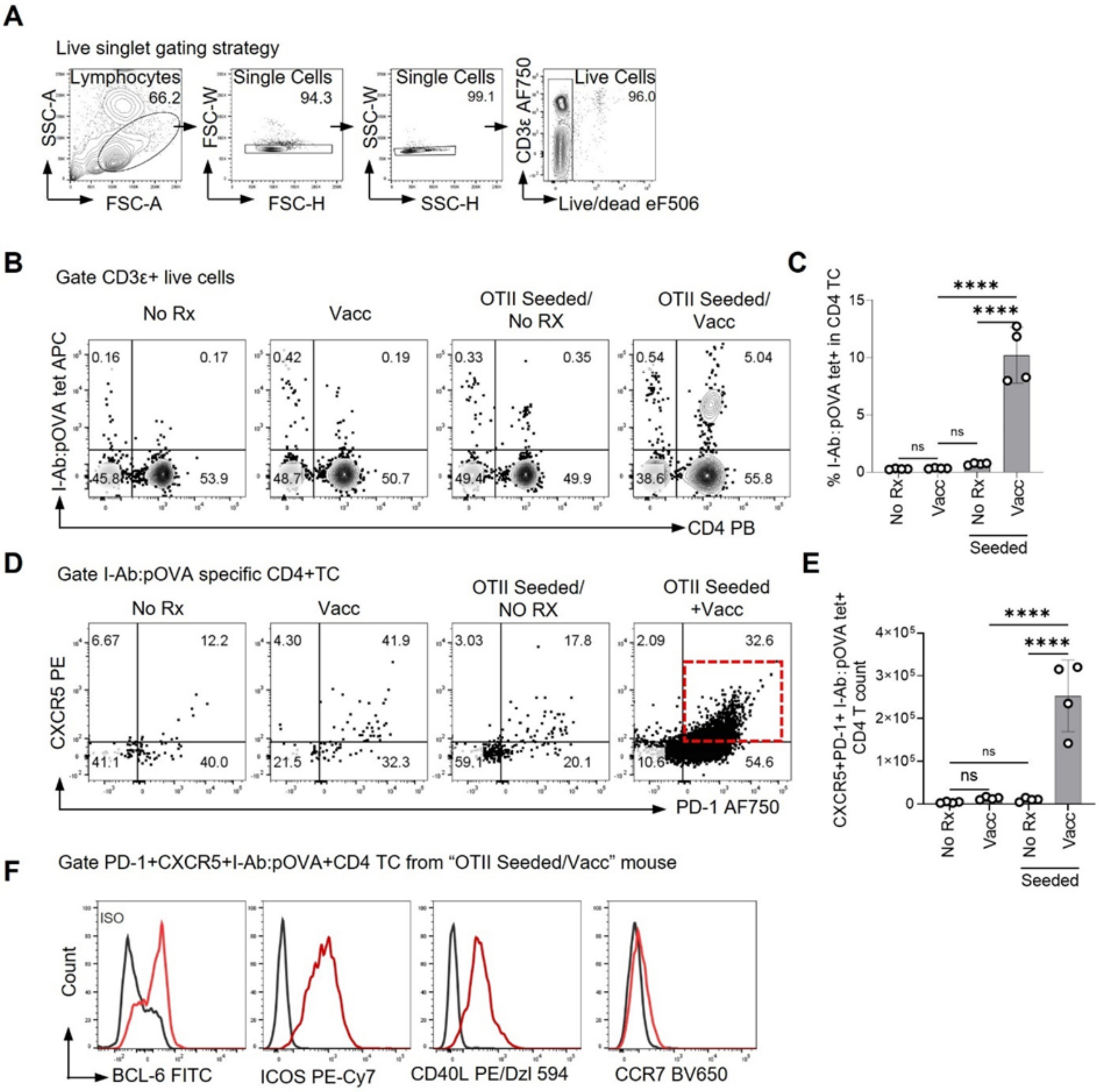


**Fig. S7. Isolation and characterization of Tfh cells from OVA-vaccinated C57bl/6 mice.** (**A**) Contour plots showing live cell and singlet gating strategy. (**B**) Contour plots showing pOVA/I-A$^b$ tetramer binding and anti-CD4 antibody staining of live T cells from unvaccinated (No Rx) and vaccinated (Vacc) C57bl/6 mice, or mice seeded OT-II T cells (OTII seeded). (**C**) Quantification of percentage of pOVA/I-Ab specific CD4$^+$ T cells 7 days post-vaccination. (**D**) Dot plots showing Tfh surface markers PD-1 and CXCR5 on pOVA/I-A$^b$-specific CD4$^+$ T cells (gated on top-right quadrants in (B)) (**E**) Quantification of count of CXCR5$^+$PD-1$^+$ pOVA/I-Ab specific CD4 T cells 7 days post-vaccination. (**F**) Histogram overlays of indicated Tfh markers (BCL-6$^+$, ICOS$^+$, CD40L$^+$, and CCR7$^-$) in CXCR5$^+$PD-1$^+$ pOVA/I-A$^b$ specific CD4$^+$ T cells gated as indicated by the red dashed line and boxed region in (D). Iso, Isotype control antibody. Means were compared by one-way ANOVA corrected for multiple comparisons. ****, $P < 0.0001$; ns, not significant ($P > 0.05$).

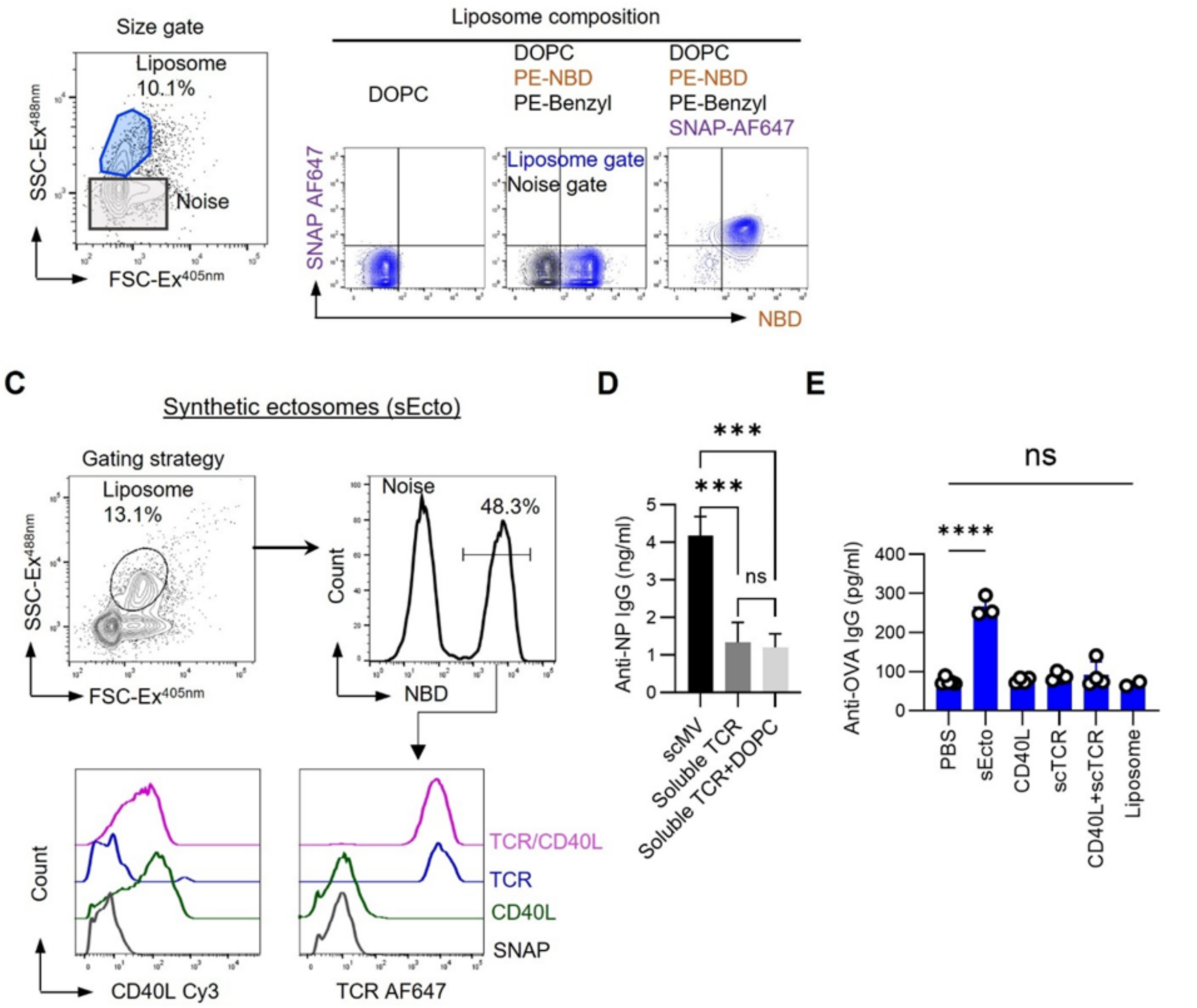


**Fig. S8. Construction, characterization and functional analysis of T cell mimetic synthetic ectosomes.** (**A**) Size gating and (**B**) contour plots of extruded liposomes composed of DOPC, DOPC with 10mol% NBD-PE, and DOPC/NBD-PE liposomes with covalently-attached AF647-labeled SNAP-tag (synthetic ectosomes – sEcto), were analyzed by small particle flow cytometry. Scatter plots show fluorescence intensity in NBD and AF647 channels with the indicated liposome compositions. (**C**) Analysis of TCR and CD40L levels (Bottom panels) on size (Left panel) and NBD fluorescence (Right-panel) serial-gated synthetic ectosomes by small particle flow cytometry. **(D)** NP-OVA pulsed QM B cells ($1\times10^5$) were incubated with sEcto ($2.7\times10^{10}$ liposomes with 1.7µg liposome-conjugated scOT-II-SNAP TCR) or the equivalent amount of soluble monomeric scOT-II-SNAP TCR (1.7µg), with or without a sMV-equivalent amount of DOPC liposomes ($2.7\times10^{10}$ liposomes). Culture supernatants were assayed for anti-NP IgG antibodies after 7 days incubation. Results are representative of 3 independent experiments. **(E)** Aged C57bl/6 mice (75-78 week-old) were immunized *s.c.* with OVA and HEL antigens and CpG ODN 2395, followed 1 hrs later by *i.v.* infusion of sEcto (12µg TCR, 12µg CD40L, $8\times10^{10}$ liposomes) or 12µg of soluble scOT-II-SNAP TCR and CD40L-SNAP, either alone or together, or with $8\times10^{10}$ bare liposomes. Serum was collected from mice 4 weeks post-vaccination, and anti-OVA IgG was measured by ELISA Means in were compared by one-way ANOVA corrected for multiple comparisons. ****, $P < 0.0001$; ***, $P < 0.001$; ns, not significant ($P > 0.05$).

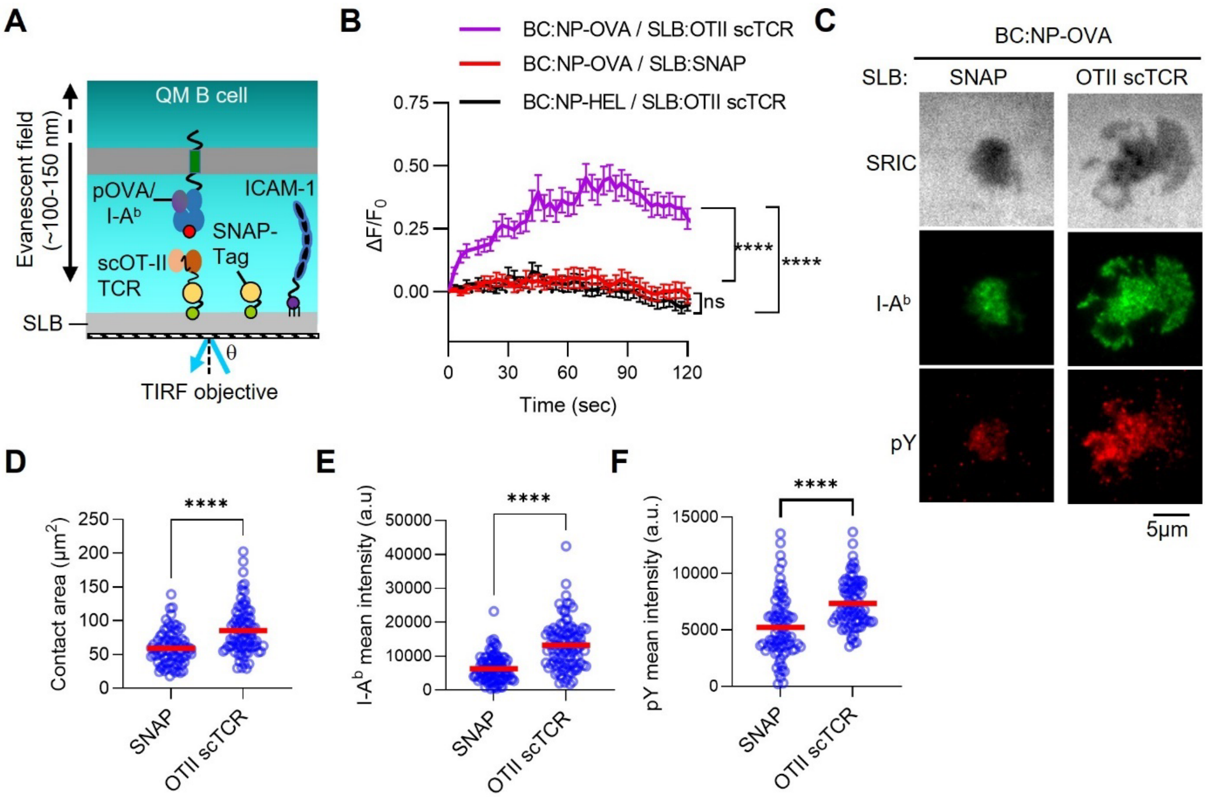


**Fig. S9. QM B cell signaling in response to TCR-mediated engagement of cognate pOVA/I-A$^b$.** (**A**) Schematic of TIRFM and supported lipid bilayer (SLB) setup for imaging QM B cell synapses. SLBs contained 200mol/µm$^2$ OT-II-SNAP or SNAP (attached to benzylguanine lipid headgroups – green circles) and 350 mol/µm$^2$ ICAM-1-10His (attached to $Ni^{3+}$/NTA lipid headgroups – purple circles). Antigen-pulsed B cells were washed and settled onto bilayers for imaging. B cells were loaded with Fluo-4 for intracellular $Ca^{2+}$-imaging by epifluorescence microscopy. For TIRFM imaging of B cell synapses, cells were incubated on SLBs containing OT-II-SNAP or SNAP for 10min at 37°C, fixed and labeled with antibodies against I-A$^b$ and phosphoptyrosine (pY). SRIC images were acquired to define synaptic contact area. (**B**) Intracellular $Ca^{2+}$-flux of B cells pulsed for 24hrs with NP-OVA or NP-HEL and IL-4, loaded with Fluo-4, and imaged on bilayers. Images were acquired every 3s and images quantitated for Fluo-4 fluorescence from first B cell contact with bilayers ($F_0$). T=0 was determined as the first appearance of Fluo-4 fluorescence at the SLB focal plane. Measurements are presented as $\Delta F/F_0$. Data are mean ± s.d., pooled from 3 independent experiments. $n$=129 cells for BC:NP-OVA/SLB:OTII scTCR, $n$=129 cells for BC:NP-OVA/SLB:SNAP, $n$=160 cells for BC:NP-HEL/SLB OTII scTCR. Mean $\Delta F/F_0$ at T=120s was compared by one-way ANOVA, corrected for multiple comparisons. (**C**) TIRFM images of NP-OVA-pulsed QM B cell synapses on SLBs containing OT-II-SNAP ($n$=85) or SNAP ($n$=81) and ICAM-1. Fixed and permeabilized cells were labelled with AF488-conjugated anti-I-A$^b$ antibody, and a pY-specific rabbit antibody followed by AF647-conjugated F(ab')$_2$ secondary antibody fragment. (**D**)-(**F**) Quantitation of B cell synapse contact area as imaged by SRIC (**D**), I-A$^b$ intensity at synaptic contacts (**E**), and pY fluorescence intensity at synaptic contacts (**F**), Data points represents individual cells. Data were representative of 3 independent experiments; a.u., arbitrary units. Means were compared by two-tailed $t$-test. ****, $P < 0.0001$; ns, not significant ($P > 0.05$).

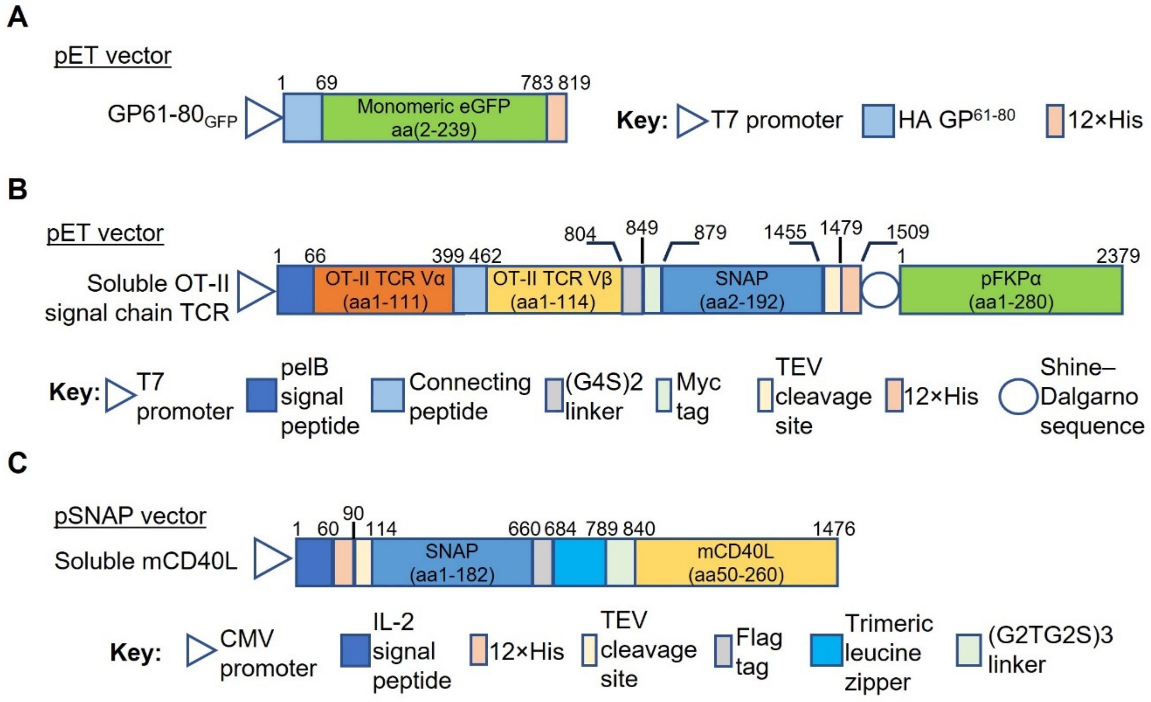


**Fig. S10. Expression constructs and vector maps.** Construct and vector details for (**A**) GFP-GP$^{61\text{-}80}$ fusion protein (for activation of GP$^{61\text{-}80}$-specific SMARTA TCR transgenic T cells), (**B**) SNAP-tag fusions of mouse OT-II and human HA1.7 single-chain TCR V domains, (**C**) SNAP-tag fusion with mouse CD40L.

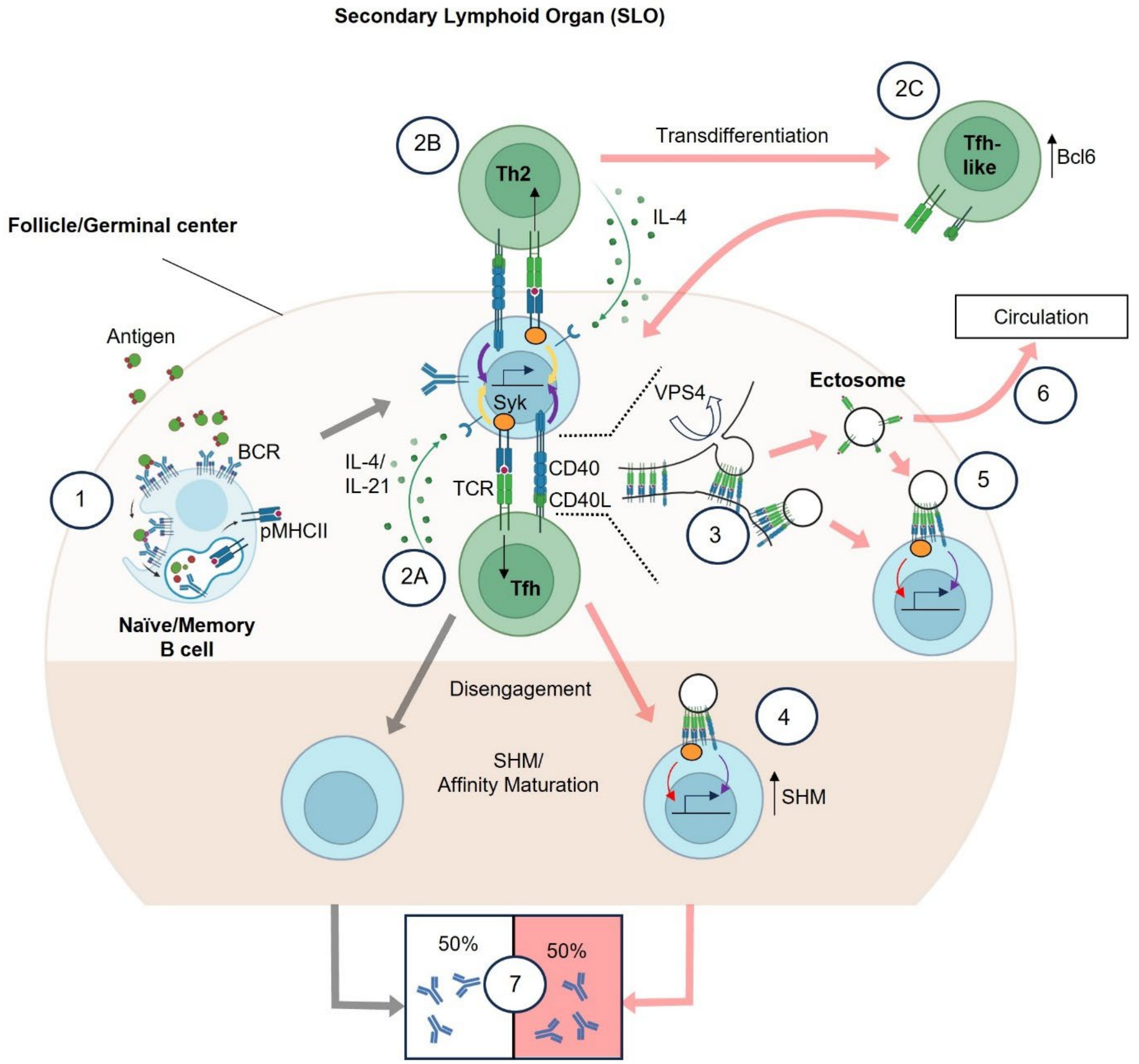


**Fig. S11. Model of T cell ectosome-mediated help for B cell antibody production.** (**1**) Antigen-specific follicular B cells encounter exogenous antigens which engage cognate surface-expressed BCRs, activating them to internalize, process and present antigen-derived peptides complexed with MHCII (pMHCII). (**2A**) Antigen-specific follicular/germinal center Tfh cells or (**2B**) extrafollicular Th2 cells at the follicular boundary, recognize and are activated by B cell-presented pMHCII at T-B cell contacts. (**2C**) Plasticity in T helper cell differentiation states may also enable Th2 transdifferentiation to a Tfh-like phenotype, permitting follicular entry and access to germinal center B cells. Activated T helper cells upregulate CD40L and helper cytokine expression which provide conventional contact-dependent and soluble help to B cells (gray arrows). (**3**) Our findings (light pink arrows) show that T helper cells also release ectosomes, which are richly decorated with TCR and CD40L, along with numerous adhesion receptors, such as SLAM family members and integrins, that promote avid interactions with B cells. T cell synaptic TCRs engage cognate pMHCII antigens on B cells, resulting in ESCRT-dependent budding and VPS4-driven scission, releasing ectosomes into the extracellular space, or resulting in their transfer to B cells *via* maintained avid adhesive contacts, or diffusion across the synaptic cleft. (**4**) Upon disengagement from T helper cells, T cell ectosomes remain bound to B cells and cluster pMHCII triggering intracellular signaling through Syk, resulting in intracellular $Ca^{2+}$ influx and transcriptional activation. This, along with engagement of CD40 by ectosome-expressed CD40L enables transferred ectosomes to continue providing long-lived help to B cells. (**5**) Released ectosomes may also diffuse within follicles to activate bystander antigen-presenting B cells displaying the cognate peptide epitope. (**6**) Alternatively, T cell ectosomes may exit local SLO, potentially providing long range help to antigen-presenting B cells (and dendritic cells) in anatomically distant tissues. (**7**) Ectosome interactions with B cells accounts for as much as 50% of the total help received by B cells for antibody production.

**Table S1. List of antibodies and recombinant proteins used in the study.**

| Protein | Host | Clone | Catalog | Company | Application |
|---|---|---|---|---|---|
| Anti-CD247 (CD3ζ) | Mouse | 8D3 | 51-6527GR | BD Pharmingen | WB |
| Anti-β-Actin | Rabbit | 13E5 | 4970 | Cell Signaling Technology | WB |
| Anti-β-Actin | Mouse | 8H10D10 | 3700 | Cell Signaling Technology | WB |
| Anti-TSG101 | Rabbit | EPR7130(B) | ab125011 | Abcam | WB |
| Anti-Flag tag | Rabbit | 8H8L17 | 701629 | Invitrogen | WB |
| Anti-VPS4B | Rabbit | Poly | 17673-1-AP | Proteintech | WB |
| Anti-CD3ε | Rabbit | E4T1B | 78588 | Cell Signaling Technology | WB |
| IRDye® 680LT Donkey anti-Mouse IgG | Donkey | Poly | 926-68022 | LI-COR | WB |
| IRDye® 800CW Donkey anti-Rabbit IgG | Donkey | Poly | 926-32213 | LI-COR | WB |
| Anti-phospho-Tyrosine (P-Tyr-1000) | Rabbit | MultiMab | 8954 | Cell Signaling Technology | WB/IF |
| Anti-CD16/32 | Rat | 93 | 101320 | Biolegend | FC |
| Anti-His-Tag | Mouse | 3D5 | R93025 | Invitrogen | FC |
| Alexa Fluor 647 F(ab')2-Goat anti-Rabbit IgG (H+L) | Goat | Poly | A-21237 | Invitrogen | FC/ IF |
| Alexa Fluor 647 F(ab')2-Goat anti-Mouse IgG (H+L) | Goat | Poly | A-21246 | Invitrogen | FC |
| APC anti-CD19 | Rat | 1D3/CD19 | 152410 | Biolegend | FC |
| Pacific Blue anti-CD19 | Rat | 6D5 | 115523 | Biolegend | FC |
| FITC anti-CD69 | ArHamster | H1.2F3 | 104506 | Biolegend | FC |
| BV421 anti-CD69 | ArHamster | H1.2F3 | 104527 | Biolegend | FC |
| PE anti-CD69 | ArHamster | H1.2F3 | 104508 | Biolegend | FC |
| BV421 ArHamster IgG | Arhmaster | HTK888 | 400935 | Biolegend | FC |
| Pacific Blue anti-CD3ε | Rat | 17A2 | 100214 | Biolegend | FC |
| APC/Fire 750 anti-I-Ab | Mouse | AF6-120.1 | 116424 | Biolegend | FC |
| Apc anti-I-Ab | Mouse | AF6-120.1 | 116417 | Biolegend | FC |
| PE-Cy5 anti-CD40 | Rat | 1C10 | 15-0401-81 | eBioscience | FC |
| PE anti-CD154 | ArHamster | MR1 | 106506 | Biolegend | FC |
| APC/Fire 750 anti-CD3ε | Rat | 17A2 | 100248 | Biolegend | FC |
| PE anti-CD95 (Fas) | Mouse | SA367H8 | 152608 | Biolegend | FC |
| PerCP/Cyanine5.5 anti-GL7 | Rat | GL7 | 144610 | Biolegend | FC |

| | | | | | |
|---|---|---|---|---|---|
| PerCP/Cyanine5.5 anti-CD4 | Rat | RM4-4 | 116012 | Biolegend | FC |
| Brilliant Violet 605 anti-CD45.1 | Mouse | A20 | 110738 | Biolegend | FC |
| Brilliant Violet 650 anti-CD45.2 | Mouse | 104 | 109836 | Biolegend | FC |
| PerCP/Cyanine5.5 anti-CD38 | Rat | 90 | 102722 | Biolegend | FC |
| Alexa Fluor 700 anti-CD23 | Rat | B3B4 | 101632 | Biolegend | FC |
| APC anti-CD81 | Arhamster | Eat-2 | 104910 | Biolegend | FC |
| APC/Fire 750 anti-CD21/CD35 | Rat | 7E9 | 123434 | Biolegend | FC |
| PE anti-CD11a | Rat | M17/4 | 101107 | Biolegend | FC |
| PE anti-TCR β | Arhamster | H57-597 | 109208 | Biolegend | FC |
| BV650 anti-TCRβ | Arhamster | H57-597 | 109251 | Biolegend | FC |
| APC anti-GATA3 | Mouse | 653806 | 16E10A23 | Biolegend | FC |
| PE anti-CD180 | Rat | RP/14 | 117706 | Biolegend | FC |
| APC anti-CD27 | Arhamster | LG.3A10 | 124212 | Biolegend | FC |
| PE anti-CD274 | Rat | 10F.9G2 | 124308 | Biolegend | FC |
| APC anti-CD205 | Rat | NLDC-145 | 138205 | Biolegend | FC |
| Alexa Fluor 594 lectin PNA | N.A. | N.A. | L32459 | Invitrogen | FC |
| PE anti-CD45R/B220 | Rat | RA3-6B2 | 103208 | Biolgend | FC |
| PE anti-CXCR5 | Rat | L138D7 | 145504 | Biolgend | FC |
| FITC anti-CXCR5 | Rat | L138D7 | 145520 | Biolgend | FC |
| APC/Fire 750 anti-PD-1 | Rat | 29F.1A12 | 135240 | Biolegend | FC |
| BV421 anti-PD-1 | Rat | 29F.1A12 | 135221 | Biolegend | FC |
| FITC anti-BCL6 | Rat | 7D1 | 358514 | Biolegend | FC |
| PE/dazzle 594 anti-CD40L | Rat | SA047C3 | 157015 | Biolegend | FC |
| PE-Cy7 anti-ICOS | Rat | 7E.17G9 | 117422 | Biolegend | FC |
| BV650 anti-CCR7 | Rat | 4B12 | 120137 | Biolegend | FC |
| PE anti-IgM | Rat | RMM-1 | 406508 | Biolegend | FC |
| Apc/Fire 750 anti-IgD | Rat | 11-26c.2a | 405744 | Biolegend | FC |
| AF568 anti-IgG Fab2 | Goat | Poly | A-11019 | Invitrogen | FC |
| Fixable Viability Dye eFluor™ 506 | N.A. | N.A. | 65-0866-14 | Invitrogen | FC |
| FITC anti-human CD19 | Mouse | Hib19 | 302206 | Biolegend | FC |
| APC/Fire 750 anti-human CD3 | Mouse | Sk7 | 344840 | Biolegend | FC |
| APC-tetramer I-$A^b$ $OVA^{329-337}$ | N.A. | N.A. | N.A. | NIH tetramer Core | FC |

| | | | | | |
|---|---|---|---|---|---|
| APC-tetramer I-$A^b$ $CLIP^{87-101}$ | N.A. | N.A. | 59666 | NIH tetramer Core | FC |
| Alexa Fluor 647 ArHamster | ArHamster | HTK888 | 400924 | Biolegend | FC |
| APC/Fire 750 Rat IgG2b, κ | Rat | RTK4530 | 400670 | Biolegend | FC |
| APC/Fire 750 Rat IgG2a, κ | Rat | RTK2758 | 400568 | Biolegend | FC |
| PE Mouse IgG1, κ | Mouse | MOPC-21 | 400112 | Biolegend | FC |
| Pacific Blue Rat IgG2b, κ | Rat | RTK4530 | 400627 | Biolegend | FC |
| PerCP/Cyanine5.5 Rat IgG2a, κ | Rat | RTK2758 | 400532 | Biolegend | FC |
| APC Armenian Hamster IgG | Arhamster | HTK888 | 400912 | Biolegend | FC |
| PE-Cy5 Rag IgG2a | Rat | eBR2a | 15-4321-80 | eBioscience | FC |
| PE Rat IgG2a, κ | Rat | RTK2758 | 400508 | Biolegend | FC |
| PE Armenian Hamster IgG | Arhamster | HTK888 | 400907 | Biolegend | FC |
| BV650 Armenian Hamster IgG | Arhamster | HTK888 | 400945 | Biolegend | FC |
| APC Mouse IgG2b, κ | Mouse | MPC-11 | 400319 | Biolegend | FC |
| Brilliant Violet 605 Mouse IgG1, κ | Mouse | MOPC-21 | 400161 | Biolegend | FC |
| PE Rat IgG2b, κ | Rat | RTK4530 | 400607 | Biolegend | FC |
| FITC Mouse IgG1 | Mouse | MOPC-21 | 400108 | Biolegend | FC |
| APC/Fire 750 Mouse IgG1 | Mouse | MOPC-21 | 400196 | Biolegend | FC |
| Alexa Fluor 700 Rat IgG2a, κ | Rat | RTK2758 | 400528 | Biolegend | FC |
| Alexa Fluor 647 Rat IgG2b | Rat | RTK4530 | 400626 | Biolegend | FC |
| APC Rat IgG2a, κ | Rat | RTK2758 | 400512 | Biolegend | FC |
| Pacific Blue Rat IgG2b, κ | Rat | RTK4530 | 400627 | Biolegend | FC |
| Alexa Fluor 488 Mouse IgG2a, κ | Mouse | MOPC-173 | 400233 | Biolegend | FC |
| Alexa Fluor 647 Mouse IgG2a, κ | Mouse | MOPC-173 | 400234 | Biolegend | FC |
| APC/Fire 750 Mouse IgG2a, κ | Mouse | MOPC-173 | 400284 | Biolegend | FC |
| PE/Dazzle 594 Rat IgG2a | Rat | RTK2758 | 400558 | Biolegend | FC |
| Alexa Fluor 568 anti-Rat IgG2b, κ, in-house conjugated, F/P=9.2 | Rat | RTK4530 | 400601 | Biolegend | FC |

| | | | | | |
|---|---|---|---|---|---|
| Alexa Fluor 647anti-CD3ε, in-house conjugated, F/P=5.6, or 12 | Rat | 17A2 | 100253 | Biolegend | FC/SIM |
| Alexa Fluor 488 anti-I-Ab, in-house conjugated, F/P=3.8 | Mouse | AF6-120.1 | 116402 | Biolegend | FC/SIM/IF |
| Alexa Fluor 568 anti-CD3ε, in-house conjugated, F/P=10 | Rat | 17A2 | 100253 | Biolegend | FC |
| Alexa Fluor 647 anti-I-Ab, in-house conjugated, F/P=1.5 | Mouse | AF-120.1 | 116402 | Biolegend | FC |
| Cy3 anti-CD154, in-house conjugated, F/P=6.9 | ArHamster | MR1 | 106516 | Biolegend | FC |
| Cy3 ArHamster, in-house conjugated F/P=8.1 | ArHamster | HTK888 | 400901 | Biolegend | FC |
| Alexa Fluor 647 Anti-TCR β, in-house conjugated, F/P=7.1 | ArHamster | H57-597 | 109254 | Biolegend | FC |
| Quantum™ Alexa Fluor® 647 MESF | N.A. | N.A. | 647A | Bangs Laboratories | FC |
| Biotin anti-CD3ε | Rat | 17A2 | 100243 | Biolegend | IP |
| Biotin Rat IgG2b, κ | Rat | RTK4530 | 400604 | Biolegend | IP |
| Anti-IL-12/IL-23 p40 | Rat | C17.8 | 505309 | Biolegend | Acti |
| Anti-IFN-γ | Rat | XMG1.2 | 505847 | Biolegend | Acti |
| Anti-CD3ε | ArHamster | 145-2C11 | 100359 | Biolegend | Acti |
| Anti-CD28 | SyHamster | 37.51 | 102121 | Biolegend | Acti |
| Anti-I-Ab | Mouse | AF6-120.1 | 116402 | Biolegend | Acti |
| Monomer I-Ab OVA$^{329-337}$ (OVA-3) | N.A. | N.A. | N.A. | NIH tetramer Core | Acti |
| CD80 extracellular domain | Mouse | N.A. | 50446-M08H-B | Sino Biological | Acti |
| ICAM-1 extracellular domain His Tag | Mouse | N.A. | 50440-M08H | Sino Biological | SLB |
| Anti-IL-2 | Rat | JES6-1A12 | 503702 | Biolegend | ELISA |
| Biotin anti-IL-2 | Rat | JES6-5H4 | 503804 | Biolegend | ELISA |
| Anti-IL-4 | Rat | 11B11 | 504102 | Biolegend | ELISA |
| Biotin anti-IL-4 | Rat | BVD6-24G2 | 504202 | Biolegend | ELISA |
| Anti-IgM mAb (MT6A3) | Rat | MT6A3 | 3885-3-250 | MABTECH | ELISA |
| HRP-anti-IgM | Goat | Poly | PA1-84383 | Invitrogen | ELISA |
| Anti-Mouse IgG (H+L) | Goat | RMG07 | SA5-10212 | Invitrogen | ELISA |
| HRP-Anti-IgG (H+L) | Goat | Poly | 62-6520 | Invitrogen | ELISA |

| | | | | | |
|---|---|---|---|---|---|
| Anti-OVA IgG | Mouse | TOSGA A1 | 520401 | Biolegend | ELISA standard |
| Anti-OVA IgM | Mouse | 2H11A 8 | 7097 | Chondrex | ELISA standard |
| Anti-HEL IgG | Mouse | 5E8 | Ab24307 3 | Abcam | ELISA standard |
| Mouse IgG | Mouse | Poly | 02-6502 | Invitrogen | ELISA standard |
| Mouse IgM | Mouse | Poly | 02-6800 | Invitrogen | ELISA Standard |
| Recombinant mouse IL2 | N.A. | N.A. | 575409 | Biolegend | ELISA standard |
| Recombinant mouse IFN-γ | N.A. | N.A. | 575309 | Biolegend | ELISA standard |
| Recombinant mouse IL4 | N.A. | N.A. | 575609 | Biolegend | ELISA Standard |

1. All antibody unless specified all react with mouse species.
2. WB, western blot; IF, immune-fluorescence; SIM, structured illumination microscopy; FC, Flow-cytometry; Acti, Activation; SLB, supported lipid bilayer.

**Table S2.** **gRNA Sequences**

| Target | gRNA (5’-3’) |
|---|---|
| VPS4B | TCCTATTATTTGGACCACC |
| | CGGGCATGGGCTTCGGGCAA |
| | TTTGGTCGCTCTTAACAA |
| TSG101 | GCGGCGGGGAGTCATAGACC |
| | TTAGCCACGGTCAGAGTTGC |
| | AGATTCCACTTGGCATGCC |
| Non-targeting | GTATTACTGATATTGGTGGG |

**Movie S1.** Electron cryotomography reconstruction of Th2 T cell derived ectosomes. Passes show raw charge density through slices of the reconstructed tomogram (first pass) and isosurface renderings (red) of ectosome membranes (second pass). Scale bar, 100nm.
